# Molecular insights into phosphorylation-induced allosteric conformational changes in β_2_-adrenergic receptor

**DOI:** 10.1101/2021.10.01.462841

**Authors:** Midhun K. Madhu, Annesha Debroy, Rajesh K. Murarka

## Abstract

The large conformational flexibility of G protein-coupled receptors (GPCRs) has been a puzzle in structural and pharmacological studies for the past few decades. Apart from structural rearrangements induced by ligands, enzymatic phosphorylations by GPCR kinases (GRKs) at the carboxy-terminal tail (C-tail) of a GPCR also makes conformational alterations to the transmembrane helices and facilitates the binding of one of its transducer proteins named β-arrestin. Phosphorylation-induced conformational transition of the receptor that causes specific binding to β-arrestin but prevents the association of other transducers such as G proteins lacks atomistic understanding and is elusive to experimental studies. Using microseconds of all-atom conventional and Gaussian accelerated molecular dynamics (GaMD) simulations, we investigate the allosteric mechanism of phosphorylation induced-conformational changes in β_2_-adrenergic receptor, a well-characterized GPCR model system. Free energy profiles reveal that the phosphorylated receptor samples a new conformational state in addition to the canonical active state corroborating with recent nuclear magnetic resonance experimental findings. The new state has a smaller intracellular cavity that is likely to accommodate β-arrestin better than G protein. Using contact map and inter-residue interaction energy calculations, we found the phosphorylated C-tail adheres to the cytosolic surface of the transmembrane domain of the receptor. Transfer entropy calculations show that the C-tail residues drive the correlated motions of TM residues, and the allosteric signal is relayed via several residues at the cytosolic surface. Our results also illustrate how the redistribution of inter-residue nonbonding interaction couples with the allosteric communication from the phosphorylated C-tail to the transmembrane. Atomistic insight into phosphorylation-induced β-arrestin specific conformation is therapeutically important to design drugs with higher efficacy and fewer side effects. Our results therefore open novel opportunities to fine-tune β-arrestin bias in GPCR signaling.

## Introduction

G protein-coupled receptors (GPCRs) are vital seven transmembrane receptor proteins that transduce signals from the extracellular to the intracellular region of eukaryotic cells. The family of these receptors is targeted by more than 30% of currently marketed drugs for the treatment of various ailments such as cardiovascular and metabolic diseases, central nervous system disorders, inflammation, and cancer.^1-6^ GPCRs recognizes and are activated by a wide variety of extracellular stimuli, including hormones, neurotransmitters, ions, lipids, and drug molecules to form stable complexes with the intracellular signal transducers such as G proteins and β-arrestins, thereby initiating multiple downstream cellular responses.^7, 8^ In a typical GPCR activation, binding of an agonist at the extracellular orthosteric pocket causes opening of the intracellular cavity of the receptor to which a G protein binds. Such allosteric communication from the exterior to the interior via GPCRs initiates G protein-mediated signaling inside a cell.^9-11^ Subsequently, the activated receptors are phosphorylated at their cytosolic carboxy-terminus (C-tail) by G protein-coupled receptor kinases (GRKs) and enable β-arrestin engagement,^12, 13^ whereby β-arrestins block G protein signaling and initiate their own signaling pathways.^14^

The two non-visual arrestins, β-arrestin 1 and β-arrestin 2, non-specifically interact with hundreds of GPCRs to elicit a multitude of cellular responses.^15^ This implies the presence of conserved structural motifs in GPCRs that enables β-arrestin engagement at the intracellular cavity. Further, arrestin usually binds rapidly to GPCRs, which indicates that a phosphorylated receptor might be sampling arrestin-favoring conformations in its conformational ensemble frequently.^16, 17^

Recent structural studies on neurotensin receptor 1 (NTSR1), M2 muscarinic receptor (M2R), and β_1_-adrenoceptor (β_1_AR) bound to β-arrestin 1 showed, besides the phosphorylated C-tail, the intracellular loops and transmembrane (TM) core of the receptors are also involved in the GPCR-β-arrestin complex formation.^18-20^ In particular, inward movements of cytoplasmic ends of transmembrane helices 5 and 6 (TM5 and TM6) were observed in the structure of β_1_AR complexed with β-arrestin 1.^18^ Although β_2_-adrenergic receptor (β_2_AR) is a pharmacologically important and well-established model system for GPCR studies, no experimental structures are available in its phosphorylated or β-arrestin bound states to understand the phosphorylation-induced conformational changes. However, recent NMR experiments on β_2_AR reveal that the interactions of the phosphorylated C-tail with the cytosolic surface of the receptor induces an inward movement of the intracellular end of TM6, thereby altering the conformation of the transducer binding cavity. Also, this TM6 movement is coupled to an increase in separation of a methionine residue (M215^5.54^) in TM5 from a phenylalanine residue (F282^6.44^) in TM6 (superscript of amino acid residues denotes the Ballesteros-Weinstein numbering scheme of GPCRs^21^). This separation occurs sufficiently far away from the cytosolic surface of the receptor, accounting for an allosteric communication from its phosphorylated C-tail to the TM domain. In fact, the phosphorylation-induced conformational changes in TM5 and TM6 are suggested to facilitate the binding of arrestin to β_2_AR.^22^ Thus, delineating the mechanisms of allosteric communication upon phosphorylation is intriguing, and this information is vital in revealing the complete picture of β-arrestin mediated signaling in GPCRs. Further, the obtained atomistic insights will help in the rational design of drugs that selectively target GPCR signalling via either G proteins or arrestins, thereby minimizing the undesired side effects.

In this study, we have performed large-scale all-atom molecular dynamics (MD) simulations on phosphorylated and unphosphorylated receptor systems to investigate the atomistic mechanism of phosphorylation-induced conformational changes in β_2_AR. We examined the allosteric effect of C-tail residue phosphorylations by GRK2, the major G protein receptor kinase of β_2_AR. By determining the directed information transfer (transfer entropy) between residues and reorganization of inter-residue energy network upon phosphorylation, we identified the allosteric communication paths from C-tail to the transmembrane domain of the receptor. For better sampling of conformation space of the membrane-receptor system, we used Gaussian accelerated MD (GaMD), an efficient unconstrained enhanced sampling technique,^23, 24^ in addition to the conventional MD (cMD) simulations.

### Computational Methods

#### System preparation and molecular dynamics simulations

Active state X-ray crystallographic structure of β_2_AR bound to full agonist BI-167107 and G protein^25^ (PDB 3SN6) was considered for all-atom MD simulations. T4 lysozyme was removed from the chain R of 3SN6, and missing parts in the receptor were modeled using MODELLER v9.17^26^, except for the N-terminus that contains 28 residues. Three engineered mutations in the crystal structure were reverted to maintain the wild type sequence (T96^2.67^M, T98^EL1^M, and E187^EL3^N). Missing residues of extracellular loop 2 (EL2) were modeled from an antagonist bound X-ray structure^27^ (PDB 2RH1), and the missing part of intracellular loop 3 (IL3) (F240^IL3^ to S261^IL3^) and three residues at the intracellular tip of TM6 (S262^6.24^ to F264^6.26^) were modeled using the secondary structure elements predicted by Jpred 4 and PSIPRED.^28, 29^ Being an unstructured region, the C-tail of β_2_AR is missing from most of the solved structures;^30^ 72 residue long C-tail (L342 – L413) was modeled from the primary sequence also using MODELLER v9.17. The modeled C-tail structure was further optimized using a constant NVT MD simulation for 500 ns to obtain a more viable starting conformation. All the solvent exposed titrable residues were kept in their dominant protonation states and residues D79^2.50^, E122^3.41^, D130^3.49^, and H172^4.64^ were protonated.^31^ Crystallographically resolved disulfide bonds between residues, C106^3.25^ - C191^EL2^ and C184 ^EL2^ - C190 ^EL2^ were retained. The parameters for ligand BI-167107 were obtained from CHARMM General Force Field (CGenFF) ParamChem server.^32, 33^ The modeled β_2_AR structure was inserted into 1-palmitoyl-2-oleoyl-sn-glycero-3-phosphocholine (POPC) lipid bilayer and the resulting system was solvated by explicit water molecules using CHARMM-GUI membrane builder module.^34, 35^ Sodium and chloride ions were added to neutralize the system and obtain a physiological salt concentration of 150 mM. The final system in a rectangular simulation box contained 116977 atoms, including 258 lipid molecules, 25363 water molecules, 62 Na^+^ ions and 65 Cl^−^ ions and initially measured roughly 101×101×143 Å^3^ in volume.

The CHARMM36m force-field parameters were used for the receptor,^36^ CHARMM36 force-field for the lipid and ions,^37^ and the CHARMM-modified TIP3P water model^38^ was used to solvate the system. All simulations were performed using particle-mesh Ewald molecular dynamics (PMEMD) implemented in AMBER18 on graphical processing units (GPUs). The energy minimization involved 5000 steps of steepest descent followed by 5000 steps of conjugate gradient. The system with positional restraints on all heavy atoms of the protein and phosphorous atoms of lipids (with a force constant of 10.0 kcal mol^-1^ Å^-2^) and restraints on dihedral angles of lipids (250 kcal mol^-1^ rad^-2^) was slowly heated to 310 K and equilibrated in NVT ensemble for 300 ps. The system was further equilibrated in NPT ensemble and the dihedral restraints on lipid were tapered off in a stepwise manner with the restraint values 100, 50, and 25 kcal mol^-1^ rad^-2^, and subsequently, the positional restraints were tapered off by 2.5 kcal mol^-1^ Å^-2^, running 1 ns of simulation at each step until no restraint remains on the lipid bilayer. The system with restraint on protein alone was further equilibrated for 15 ns. Restraint on protein heavy atoms was removed slowly by reducing it step by step, from 10.0 kcal mol^-1^ Å^-2^ to 0, equilibrating the system for 2 ns at each of the steps with harmonic force constants 10, 7.5, 5.0, 4.0, 3.0, 2.0, 1.0, 0.8, 0.6, 0.4, 0.2, 0.1 kcal mol^-1^ Å^-2^. Langevin thermostat was used to maintain the 310K and Monte Carlo barostat was used to keep the system at 1 bar pressure throughout the simulations. The SHAKE algorithm was used for constraining bonds involving hydrogen atoms and the SETTLE algorithm was used for water molecules. Periodic boundary conditions were applied, and the cut-off for nonbonded interactions was set at 9 Å. The particle mesh Ewald (PME) method was used to calculate the long-range electrostatic interactions with Ewald coefficient of approximately 0.31 Å and a cubic B-spline interpolation of order 4. Further the system was equilibrated for 100 ns and the last frame was extracted for *in silico* phosphorylation.

Phosphorylation by GRK2 of serine and threonine residues in C-tail of the receptor (T360, S364, S396, S401, S407, and S411)^39, 40^ was performed using CHARMM-GUI server. To attain local equilibration around the newly added phosphate groups, we ran cMD simulations for 25 ns after each phosphate group addition. Lastly, we performed an additional 100 ns equilibration simulation of the phosphorylated system in NPT ensemble.

##### Gaussian accelerated molecular dynamics (GaMD) simulations

To alleviate the poor sampling of the conformational space of biomolecular systems of large size in cMD, enhanced sampling simulation techniques are routinely employed.^41-43^ Here we used the recently developed GaMD technique that achieves unconstrained enhanced sampling of biomolecules. GaMD has been successfully demonstrated in multiple studies of class A GPCR systems.^44-46^ It is shown to accelerate the conformational transitions of proteins by adding a harmonic boost potential, *ΔV*(**r**), to the system potential, *V*(**r**), to overcome the energy barrier,^23, 24^

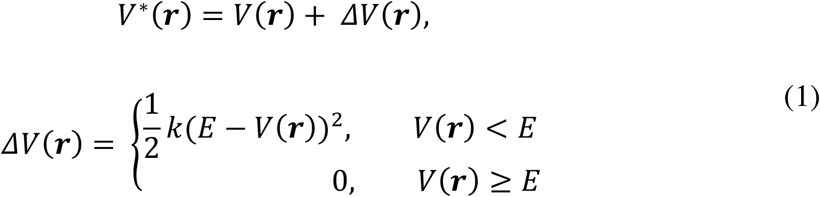

where *V*^*\**^(***r***) is the modified system potential, ***r***= {***r*_1_, *r*_2_**, **…**, ***r*_*N*_**} is the coordinate vector of *N* atoms in the system, *k* is the harmonic force constant, and *E* is the threshold cut-off for *V*(**r**), below which the boost potential is added.

The GaMD simulations of the unphosphorylated and phosphorylated receptors (hereafter referred to as B2AR and B2ARP, respectively) were performed using the GaMD module of PMEMD implemented in AMBER18 software on GPUs.^40, 47^ We used the standard GaMD simulation protocol as described in previous simulation studies on GPCRs.^44-46^ Starting from the last frame of the previous equilibration step of each system, a 12 ns short cMD simulation was run to collect potential statistics, e.g., the maximum (*V*_max_), minimum (*V*_min_), average (*V*_av_), and standard deviation (σ_V_) of system potential energies. After adding the boost potential, a 40 ns GaMD equilibration was performed. We then randomized the initial atomic velocities for production simulations and started three independent trajectories of 2 μs each for both B2AR and B2ARP systems. The first 100 ns of each run was further considered as equilibration and disregarded in the final analyses. GaMD simulations were run using a time step of 2 fs, and the trajectory frames were written at every 1 ps. We used “dual boost” level (applying boost potential to the dihedral energy term and the total potential energy term) of GaMD, setting the reference energy to the lower bound, i.e., *E* = *V*_max_. We calculated the average and standard deviation of boost potentials in every 800 ps, and σ_0_ (the upper limit of standard deviation) was set to 6.0 kcal/mol for both potential and dihedral energetic terms. The average and standard deviation of boost potential for each simulation are given in table S1.

Along with B2AR and B2ARP systems, we also performed another set of GaMD simulations (two independent trajectories of 1 *μ*s each) on a truncated β_2_AR with no C-tail and IL3 (as in the active crystal structure, PDB 3SN6) using the same parameters and simulation settings mentioned above. These simulations were conducted to examine the influence of their absence on the dynamics of β_2_AR.

For the free energy calculations, the canonical ensemble probability distribution (*p*(*A*)) along any reaction coordinate *A*(***r***) was recovered by reweighting its biased (GaMD) probability distribution *p*^*^(*A*), where ***r*** denotes the atomic positions, ***r*** = {***r*_1_, *r*_2_**, …, ***r*_*N*_**} and *N* denotes total number of atoms. Using the boost potential Δ*V(****r***) for each simulation frame, *p*(*A*) is calculated as,(Ref)

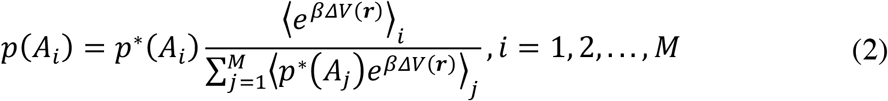

where *M* is the number of bins and (e^*βΔ*^^*V*(*r*)^)_*i*_ is the ensemble-averaged reweighting factor of *ΔV*(***r***) for simulation frames in the *i*^*t*h^ bin, where *β* = 1⁄*k*_*B*_*T* and *k*_*B*_ is the Boltzmann constant. The reweighting factor is approximated using the cumulant expansion,

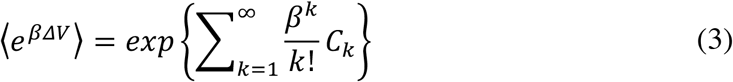

Usually *ΔV*(***r***) follows near-Gaussian distribution and thus, cumulant expansion to the second order can be a good approximation for reweighting the distribution, where the first two cumulants are given by *C*_1_ = ⟨Δ*V*⟩ and 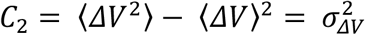. The reweighted free energy or the potential of mean force (PMF) is calculated as *F*(*A*_*i*_) = −(1⁄*β*)ln *p*(*A*_*i*_).

##### Conventional molecular dynamics (cMD) simulations

We also performed cMD simulations in NVT ensemble on both B2AR and B2ARP systems, starting from specific initial configurations from GaMD simulations (see Results and discussion section). Two independent trajectories of 500 ns length were generated for each system and coordinates were saved at every 5 ps. We used the AMBER18 package, and the simulation protocols remained the same as discussed earlier.

### Analysis of trajectories

#### Transfer entropy

Transfer entropy (TE) quantifies the directed flow of information from one random physical process *X* to another process *Y*, represented by the time series {*x*_*t*_} and {*y*_*t*_} (with discrete-valued time index *t*), respectively.^48^ Such flow of information is interpreted as the causal relationship between *X* and *Y* (influence of *X* on *Y*) and is defined as the predicted information about the future of *Y* from the past of *X* in addition to the information obtained from its own past (known as Wiener’s principle of causality). Using Taken’s time-delayed embedding^49^ to construct the state vectors of the Markov processes *X* and *Y*, the information-theoretic measure of transfer entropy from *X* to *Y* is defined as,^48, 50^

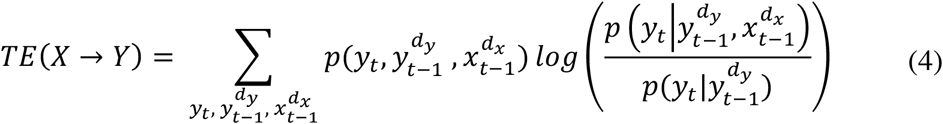

where 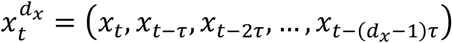 represent the delay-embedding state (delay vector) of the Markov process *X* at time *t* with embedding dimension *d*_*x*_ and embedding delay *τ*. Similarly, 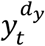 represent the process *Y* with embedding dimension 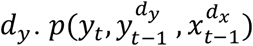 is the joint probability of observing the future state *y*_*t*_ of *Y*, and the past states 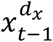 and 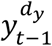 of *X* and *Y*, respectively. 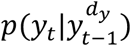 is the conditional probability to find *Y* in state *y*_*t*_, given its past 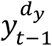, and 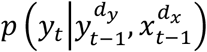 is the conditional probability to find *Y* in state *y*_*t*_, given the past of both *X* and *Y*. When the processes *X* and *Y* are independent, the future state of *Y* is not influenced by the history of *X*; the two conditional probabilities are therefore equal in this case and there is no information flow from *X* to *Y*. In fact, the transfer entropy expression (eq 4) uses the Kullback-Leibler divergence to measure the deviation between the two conditional probabilities.^48^ It is obvious that the transfer entropy is by construction asymmetric, *TE*(*X* → *Y*) ≠ *TE*(*Y* → *X*) in general.

As information transfer in physical systems is necessarily associated with a physical interaction, it always takes a finite time for information to propagate from *X* to *Y*. Thus, to take into account the true interaction delay *δ* between *X* and *Y* that maximizes the information flow, and to ensure an optimal self-prediction from the past of *Y* to the future of *Y*, eq 4 can be rewritten as,^50^

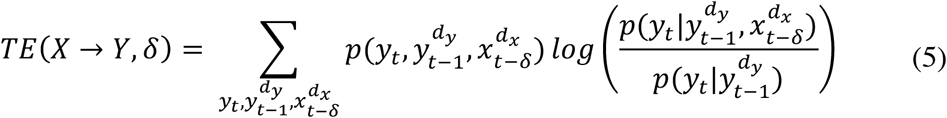

For a practical estimation of TE, eq 5 can be rewritten in terms of different joint and marginal Shannon entropies *H*,

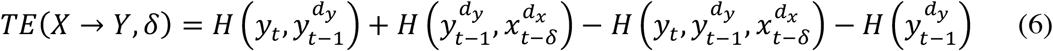

where the Shannon entropy, 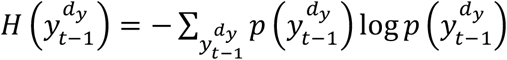 and the sum is over all states.

We considered fluctuations of alpha carbon (C_α_) of individual residues as random processes and defined the *net* entropy transfer from residue *X* to *Y* as,

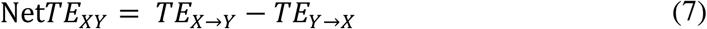

The change in Net*TE*_*XY*_ upon phosphorylation is estimated using the equation,

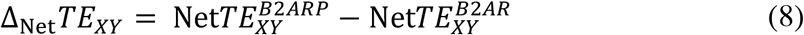

The net entropy transfer from residue *X* to all other residues can be obtained by summing the expression of Net*TE*_*XY*_ (eq 7) over all *Y*. Therefore, by summing eq 8 over all *Y*, the change in net entropy transfer from residue *X* to all other residues upon phosphorylation can be calculated as,

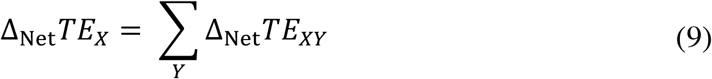

We performed n(n-1) TE calculations (where n is the number of residues in the receptor) for both B2AR and B2ARP systems using TRENTOOL Matlab toolbox.^51^ Individual entropy terms in eq 6 is calculated using the *k* nearest neighbor search method by Kraskov-Stögbauer-Grassberger^52^ on reconstructed state space (embedded time series). Embedding dimension *d* and embedding delay *τ* for the time series of each residue was optimized using Ragwitz criterion in which both are estimated jointly by minimizing the prediction error of a local predictor that predict the future of *Y* from its past (i.e., provide an optimal self-prediction).^53^ We found the values of interaction delay δ are in the range of 1.4 -1.7 ns.

Finally, the above transfer entropy estimates may give rise to false correlations between *X* and *Y* due to finite data of the two time series. For removing this finite sample bias, the TE values are compared against surrogate data sets generated by random shuffling of the original time series *X* (or *Y*) that destroys any correlations between *X* and *Y* while preserving their distributions. The TE measure thus obtained and reported in this study is the effective transfer entropy, and is given by,^54, 55^

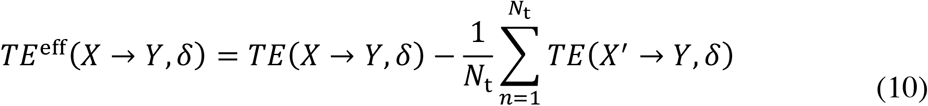

where *X*′ represents the shuffled time series of *X* and *N*_t_ is the number of times it is shuffled. For our calculations, we used *N*_t_ = 100.

#### Nonbonding interaction energies

The average nonbonding interaction energy of a residue pair is the sum of their electrostatic and van der Waals’ interaction energies,

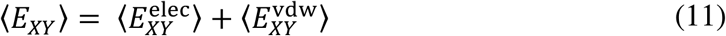

We define the change in average pairwise interaction energy upon phosphorylation as,

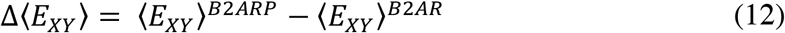

Pairwise residue interaction energies were calculated using CPPTRAJ module of AMBER18.

#### Contact maps

To identify unique contacts in the unphosphorylated and phosphorylated receptors, we computed all the contact pairs using an in-house python script. MDTraj python library was used for distance calculations.^56^ Two residues are considered to be in contact if any two of their heavy atoms are within 5 Å for more than 80% of simulation time.

Trajectories were visualized, and high-resolution pictures were rendered using the visual molecular dynamics (VMD) program.

## Results and discussion

Schematic representation of the system considered for simulations with the receptor β_2_AR inserted in a POPC bilayer, solvated in explicit water, and neutralized using 150 mM NaCl is shown in Figure S1. Summary of GaMD simulations and their average boost potentials are given in Table S1. We considered eight residues at the intracellular (cytosolic) and extracellular region of the TM helices to define their respective ends. Further, eight residues in the middle of a TM helix is considered as the central part of that helix. A detailed residue-wise description of TM helices (and other structural elements) is given in Table S2.

### Free energy landscape of the phosphorylated receptor sampled a new conformational state

We used TM3-TM6 distance and the distance between the center of mass of the side chains of M215^5.54^ and F282^6.44^ for the potential of mean force (PMF) calculations. Both are major structural variations observed in NMR spectra of phosphorylated β_2_AR; the former corresponds to the movement of the intracellular half of TM6, and the latter represents a conformational change in the mid transmembrane region of TM5 and TM6.^22^ The free energy landscapes revealed that compared to the unphosphorylated receptor (B2AR), the phosphorylated receptor (B2ARP) samples a larger conformational space (Figure 1a, b). While B2AR sampled a single free energy minimum (m_1_ in Figure 1a) closer to the G protein-bound active structure (PDB 3SN6), the free energy surface of B2ARP shows two minima, of which one (m_1_’ in Figure 1b) is similar to the minimum observed in B2AR and the other one (m_2_ in Figure 1b) represents a new conformational state. Though conformations from the minima m_1_ and m_1_’ are observed to be quite similar (Figure S2), m_2_ showed significant structural deviations (Figure 1c). In the newly sampled state, TM6 is moved inward by ∼4 Å, and the distance between residues M215^5.54^ and F282^6.44^ is increased by ∼3 Å (Figure 1d, e). These findings corroborate well with the results obtained in previous NMR experiments,^22^ suggesting that our simulations could successfully capture the phosphorylation-induced allostery in β_2_AR.

**Figure 1.**
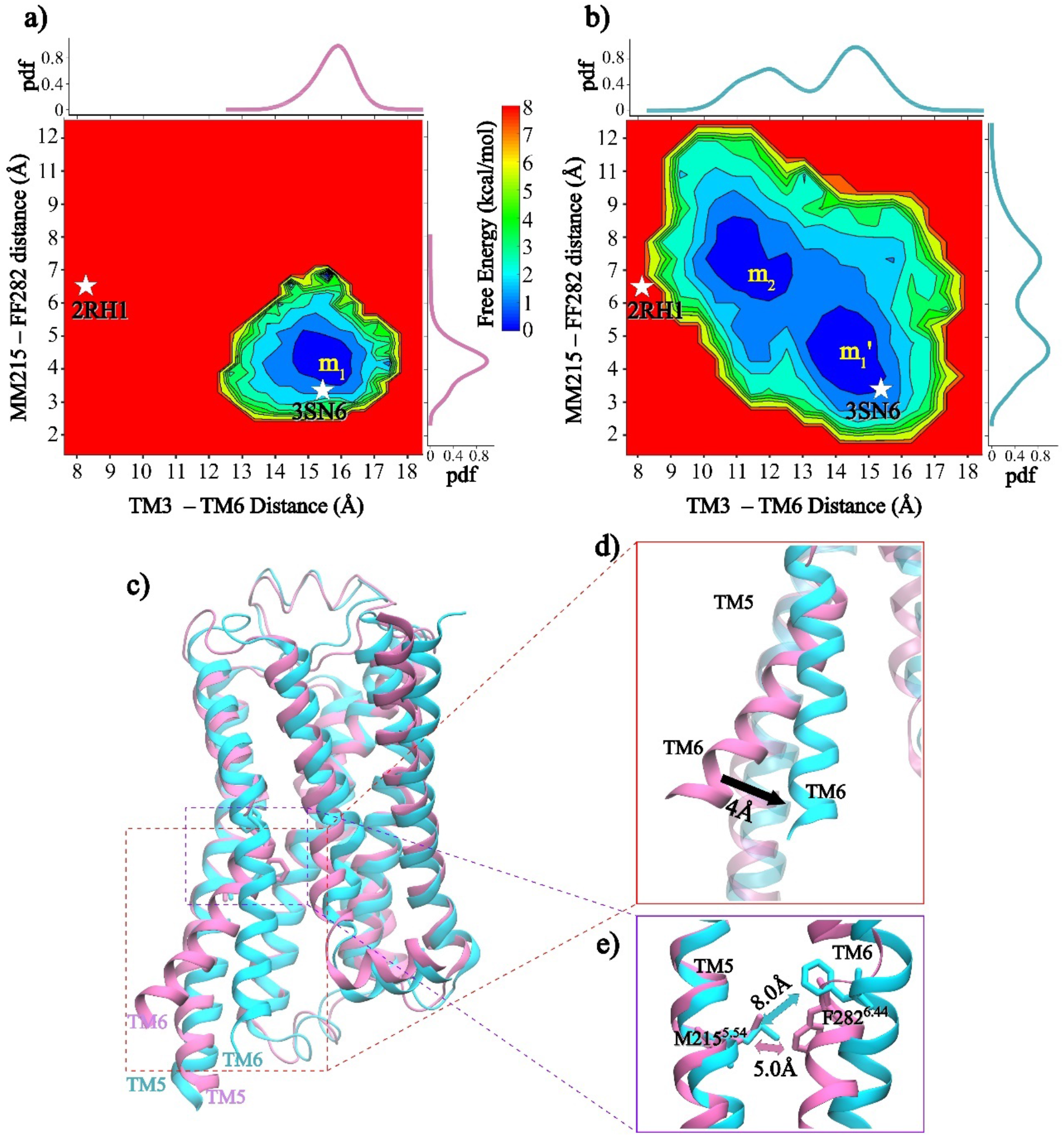
The two-dimensional free-energy landscape of **a)** unphosphorylated receptor (B2AR, pink) **b)** phosphorylated receptor (B2ARP, cyan), plotted against the collective variables, TM3-TM6 distance (distance between C_α_ of R131^3.50^ and L272^6.34^) and the center of mass distance of M215^5.54^ (TM5) and F282^6.44^ (TM6) side chains. Stars denote the active crystal structure (PDB 3SN6) and the inactive crystal structure (PDB 2RH1). The probability distribution of each collective variable is depicted on top of the free-energy plot. **c)** Representative conformations from m_1_ (pink) and m_2_ (cyan). **d)** Movement of TM6 at the cytosolic end. **e)** Separation of F282^6.44^ and M215^5.54^.

We noted that contrary to a previous large-scale unbiased MD simulation study on the agonist bound β_2_AR,^31^ our GaMD simulations did not sample the inactive state of the receptor (Figure S3). The model system used in that study was a truncated receptor with no C-tail and IL3 residues missing as in the crystal structure. For comparison, we performed GaMD simulations with a similar starting structure. Our results showed that the receptor attained an inactive state characterized by an inward movement of the intracellular end of TM6 and the conserved NPxxY^7.53^ motif (Figure S4) switches to a conformation closer to the inactive crystal structure (PDB 2RH1). In our simulations, inclusion of both C-tail and IL3 could be the reason for the agonist-bound (unphosphorylated and phosphorylated) receptors not sampling spontaneously a conformational state closer to the inactive state. In fact, it is worth mentioning that both the structural elements, C-tail and IL3, are reported to be involved in spontaneous activation of β_2_AR.^57^

### Structural variations in the transmembrane domain and change in residue-residue contacts upon phosphorylation

We compared the structures obtained from minima m_1_ and m_2_ (Figure 1a, b) to identify the structural variations in the conformational state sampled by the phosphorylated receptor (average structures taken from these minima are shown in Figure S5). On the extracellular part of the receptor, we observed an outward movement of TM5, TM6, and TM7 (∼1 Å each) in B2ARP compared to B2AR. Further, both EL2 and EL3 in B2ARP showed ∼1 Å shift from that of B2AR. Consequently, the volume of the ligand-binding pocket is increased to 482.6 ± 66.4 Å^3^ in B2ARP from 407.8 ± 51.5 Å^3^ in B2AR (Figure S6a, b). Because of such rearrangements of different extracellular helices and loops, the agonist BI-167107 lost multiple contacts in B2ARP, especially with the residue I169^4.61^ in TM4, Y174 and T195 in EL2, F208^5.47^ in TM5, and Y308^7.35^ in TM7. In addition, we observed that the agonist formed unique contacts with the residue D113^3.32^ in TM3, S207^5.46^ in TM5, and V297^6.59^ in TM6 (Figure 2). It is to be noted that the extracellular TM5 and TM6 of β_1_AR also moved outward in its β-arresin bound structure. When compared with the G protein bound structure of β_2_AR, β_1_AR bound to β-arresin 1 showed contact rearrangements with the agonist ligand,^20^ where the ligand made fewer contacts with TM3 and TM5 and additional contacts with TM6.

**Figure 2.**
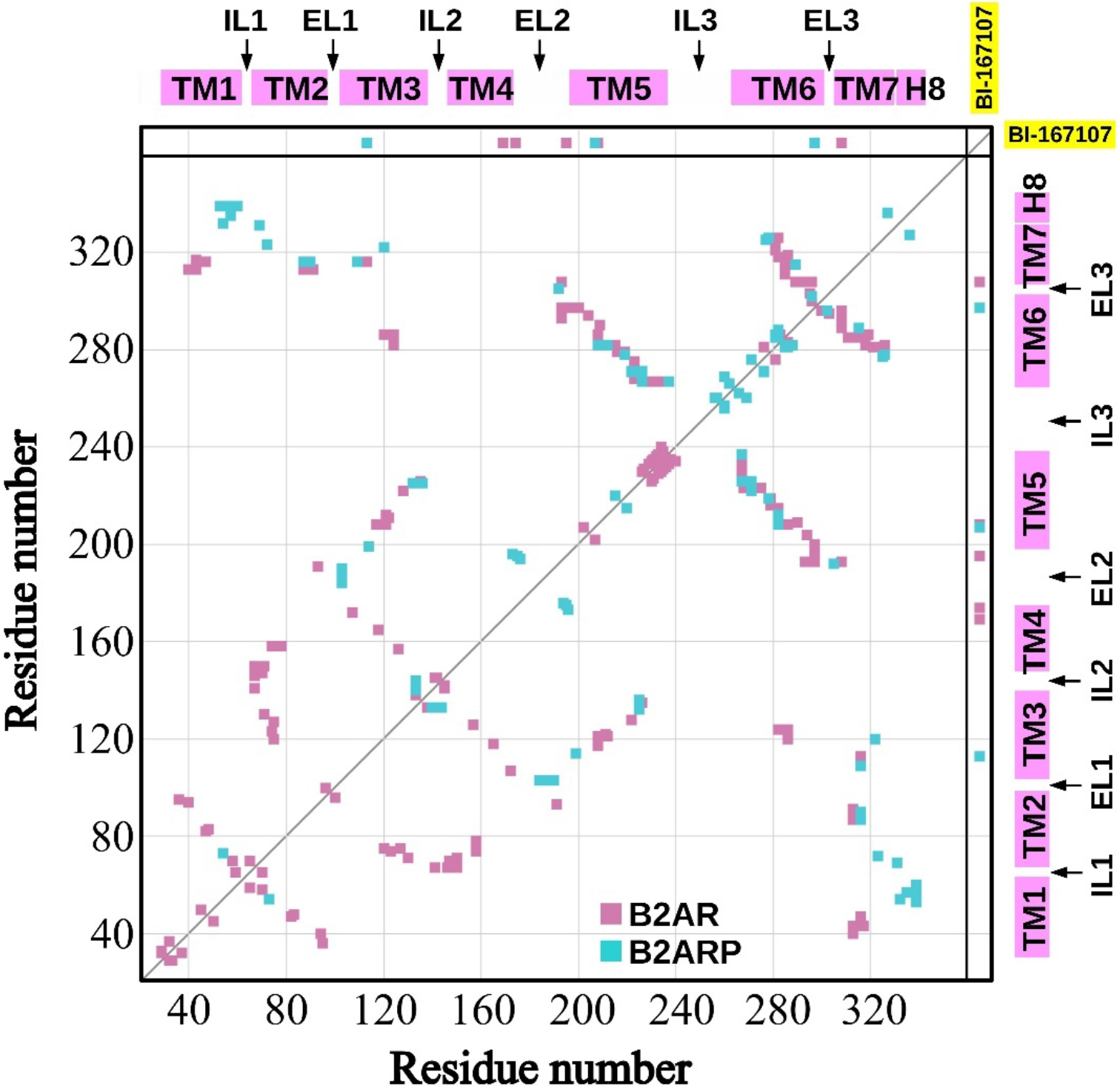
Contact map showing unique contacts formed between residues of TM region (TM helices and loops) in B2AR (pink) and B2ARP (cyan). The panel on the right represents the unique contacts formed by residues in the TM region and the agonist ligand BI-167107.

In the intracellular region, besides the significant inward motion of TM6 (∼4 Å), TM3 and TM5 moved slight inward (∼2.0 Å) and outward (∼1.5 Å), respectively, as compared to B2AR (Figure S5). The net inward motion of intracellular end of helices suggests a reduction in size of the transducer binding cavity. In fact, our calculations show that the volume of the intracellular cavity is reduced to 648.3 ± 151.1 Å^3^ from 930.6 ± 176.9 Å^3^ in B2AR (Figure S6c, d). We note that, in the cryo-EM structure of β_1_AR-β arrestin 1 complex,^20^ the intracellular ends of TM5 and TM6 of the receptor were observed to move inward, implying a decrease in the volume of the transducer binding cavity. Further, the structural study also revealed that the finger loop of β-arrestin 1 binds to a narrower cleft at the cytosolic surface of the β_1_AR as compared to the α5 helix of G protein which requires a broader binding region in the intracellular cavity of β_2_AR.^20^ Moreover, surface plasmon resonance (SPR) experiments showed that a G protein mimetic nanobody selectively recognizes the unphosphorylated β_2_AR having a larger outward movement of TM6, whereas the SPR response diminishes for the phosphorylated receptor with a relatively smaller TM6 outward movement.^22^ Thus, the reduction in the volume of the transducer binding cavity in β_2_AR upon phosphorylation observed in our simulations corroborates with the experimental findings.

The rearrangement of residue contacts plays a crucial role in allosteric communication in proteins. To identify such residue level structural variations of the receptor due to phosphorylation, we compared the unique contacts formed in B2AR and B2ARP (Figure 2, Table S3). In the TM region (TM helices and loops), the structural rearrangements in B2ARP caused breaking of several inter-residue contacts and the number of unique contacts reduced to half of that found in B2AR. Upon phosphorylation, extensive formation and breaking of contacts are observed mostly in the extracellular and intracellular ends of the helices and loops. We observed that the outward movement of extracellular ends of TM5, TM6, and TM7 caused breaking of many contacts in B2ARP; the contacts of TM3 with TM5 (e.g., I121^3.40^ with F208^5.47^ and E122^3.41^ with P211^5.50^), TM6 with TM5 (e.g., I294^6.56^ with S204^5.43^; V297^6.59^ with Q197^5.36^ and A200^5.39^), and TM6 with both EL3 and TM7 (e.g., H296^6.58^ with D300^EL3^; F289^6.51^ with Y308^7.35^) are found to be broken (Figure 2, Table S3). We also observed the formation of a number of contacts between TM3 and EL2, such as N103^3.22^ with C184^EL2^, A186^EL2^, N187^EL2^, and C190^EL2^, in the extracellular end of the receptor.

The inward movement of TM6 and the outward movement of TM5 in the intracellular end caused the breaking of several contacts (e.g., F223^5.62^ with E268^6.30^, L272^6.34^, and L275^6.37^; D234^5.73^ with E237^IL3^ - F240^IL3^ and K267^6.29^). In addition, due to the shift of IL2 (Figure S5), TM2 broke several contacts with IL2 as well as with its connecting helices TM3 and TM4, such as F71^2.42^ with D130^3.49^, V67^2.38^ with Y141^IL2^, and V67^2.38^ with T146^4.38^ (Figure 2, Table S3). Besides the contact rearrangements in the intracellular and extracellular regions, redistribution of residue contacts is also observed in the central part of different TM helices. In particular, multiple contacts between TM3 and TM5 (e.g., I121^3.40^ with F208^5.47^ and L212^5.51^; E122^3.41^ with P211^5.50^) as well as between TM5 and TM6 (e.g., M215^5.54^ with F282^6.44^; Y219^5.58^ with M279^6.41^) are broken upon phosphorylation of β_2_AR (Figure 2, Table S3). Notably, the breakage of the contacts of methionine residues M215^5.54^ and M279^6.41^ with F282^6.44^ and Y219^5.58^, respectively, is likely an implication of the allosteric conformational variations around M215^5.54^ and M279^6.41^ observed in NMR experiments.^22^

### C-tail adheres to the cytosolic surface of β_2_AR and samples a narrow conformational space upon phosphorylation

Experimental studies show that C-tail is one of the disordered motifs in β_2_AR and post-translational modifications such as phosphorylation of this motif is important in the modulation of GPCR signaling.^30^ In Figure 3a, we show the dynamics of C-tail upon phosphorylation by comparing its conformational distributions as functions of radius of gyration (*R*_*g*_) and root-mean-square deviation (RMSD) in B2AR and B2ARP (minima m_1_ and m_2_, Figure 1a, b). In B2AR, the barycenter of the scattered plot appears at *R*_*g*_ of 16.5 Å and RMSD of 10.1 Å and in B2ARP, it appears at *R*_*g*_ of 13.8 Å and RMSD of 3.1 Å. The smaller *R*_*g*_ and RMSD values in B2ARP compared with B2AR imply that the structural ensemble of C-tail samples a more stable and compact state upon phosphorylation.

**Figure 3.**
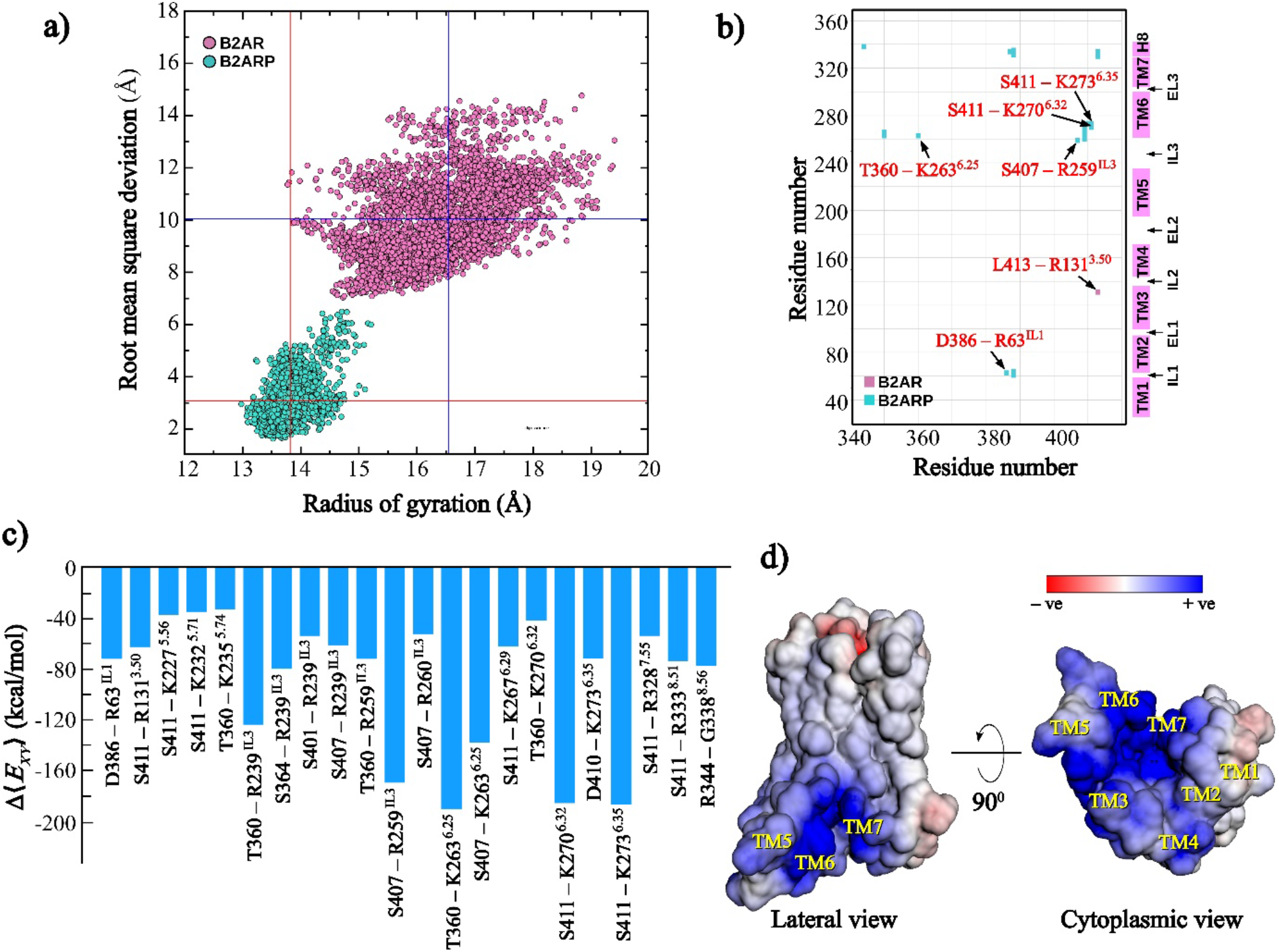
Conformational differences of C-tail in B2AR and B2ARP. **a)** Conformational distribution of C-tail as functions of *R*_*g*_ and RMSD in B2AR and B2ARP. The blue lines represent the *R*_*g*_ and RMSD values corresponding to the barycenter of the scattered plot distribution in B2AR and the red lines represent the same in B2ARP. **b)** Contact map showing the unique contacts formed between the C-tail residues and the TM residues in B2AR and B2ARP. **c)** Difference in interaction energies (|Δ⟨*E*_XY_⟩| > 30 kcal/mol) of significant residue pairs at the interface of the C-tail and the TM domain. **d)** Surface charge potentials of β_2_AR calculated using Adaptive Poisson-Boltzmann Solver (APBS).

Next, we examined the unique contacts formed by the C-tail with the TM helices and intracellular loops of the receptor. C-tail residues formed extensive contacts with the TM region in B2ARP as compared to B2AR (seventeen against only one) as shown in Figure 3b and Table S3. Mostly the residues from IL1, IL3, H8, and the intracellular ends of TM6 and TM7 formed contacts with the phosphorylated C-tail, indicating that the C-tail has a strong affinity to bind with the cytosolic surface of the receptor upon phosphorylation. Importantly, several contacting residue pairs including, (T360, K263^6.25^), (S407, K259^IL3^), (S411, K270^6.32^), and (S411, K273^6.35^), involved phosphorylated residues. These residue-residue contacts found at the interface of the intracellular region and C-tail are likely to influence the communication from the C-tail to the TM domain and therefore play a vital role in phosphorylation induce allostery in β_2_AR.

To further examine the origin of the structural stability of C-tail conformations upon phosphorylation, we calculated the change in average pairwise interaction energies Δ⟨*E*_*XY*_⟩ (eq 12) between the C-tail and TM region residues from the two conventional MD simulation trajectories (a cumulative length of 1*μ*s) that were started from the minima m_1_ and m_2_ (Figure 1a, b). It is to be noted that a negative (or positive) value of Δ⟨*E*_*XY*_⟩ signifies the interaction between residues *X* and *Y* is more (or less) stable in B2ARP than in B2AR. For the residue pairs with |Δ⟨*E*_*XY*_⟩| > 30 kcal/mol, we found that they are largely stabilizing interactions; in particular, the phosphorylated residues such as T360, S364, S407, and S411 established strong favorable interactions with positively charged residues in IL3, H8, and the intracellular ends of TM3, TM5, TM6, and TM7 (Figure 3c, Table S4). Besides the phosphorylated residues, other negatively charged residues from the C-tail such as D386 and D410 also formed stable interactions with the TM region of B2ARP. It is known that class A GPCRs possess a positively charged residue cluster on the cytosolic face of the TM region.^22, 58-60^ Surface charge potentials calculated using adaptive Poisson-Boltzmann solver (APBS)^61^ showed a similar cluster of concentrated positive charges on the cytosolic surface of β_2_AR (Figure 3d). Our simulation results are in good agreement with the previous NMR experimental findings,^22^ suggesting the adherence of the C-tail to a positively charged residue cluster in the intracellular regions of the phosphorylated β_2_AR and its involvement in the structural rearrangements of the TM domain.

### Entropy transfer reveals allosteric communications between different parts of the receptor

To unravel the allosteric communication from the phosphorylated C-tail to the TM domain of the receptor, we analyzed the correlated residue motions of all residue pairs using transfer entropy (TE) measures. It should be noted that TE estimates are quite expensive computationally, since each calculation requires to be performed in a higher dimensional delay-embedded state space (see the Computational Methods section). Thus, we considered 100 ns of well-equilibrated samples from the conventional MD simulations of both the systems starting from the minima m_1_ and m_2_ (Figure 1a, b) for TE calculations. The positive or negative sign of Net*TE*_*XY*_ between two residues *X* and *Y* calculated using eq 7 shows whether a residue acting as a source (donor) or sink (acceptor) of information transfer. The entropy sources drive the correlated motions, whereas the entropy sinks are the responders. We also note that a positive value of *Δ*_Net_*TE*_*XY*_ (eq 8) implies residue *X* becomes either of the following upon phosphorylation: (i) changes its entropy transfer characteristic from acceptor to donor, (ii) a stronger entropy donor, or (iii) a weaker entropy acceptor. Whereas, a negative value of *Δ*_Net_*TE*_*XY*_ indicates opposite characteristics of residue X.

Figure 4a, b represents Net*TE*_*XY*_ values between all residue pairs of individual systems (B2AR and B2ARP, respectively), and Figure 4c represents *Δ*_Net_*TE*_*XY*_ (subtracting the TE values in 4a from 4b). We observed that the C-tail residues upon phosphorylation mostly drive the correlated motions in different TM regions of the receptor (Figure 4b, c, zone 1). Figure 4d shows the change in net transfer entropy from each TM residue *X* to all the C-tail residues *Y* (obtained by summing over all *Y*, eq 9). We observed that the central regions of several TM helices accept entropy from the C-tail residues; specifically, TM3, TM4, TM5, and TM6 (region marked in Figure 4d) act as entropy sinks. The intracellular ends of TM1, TM2 and their connecting loop (IL1) appeared to accept entropy from C-tail and the extracellular ends of TM4, TM5, and TM6 also showed a similar trend. In addition, the helix H8 and a number of residues in the intracellular half of TM6 are found to act as entropy acceptor from C-tail (Figure 4d).

**Figure 4.**
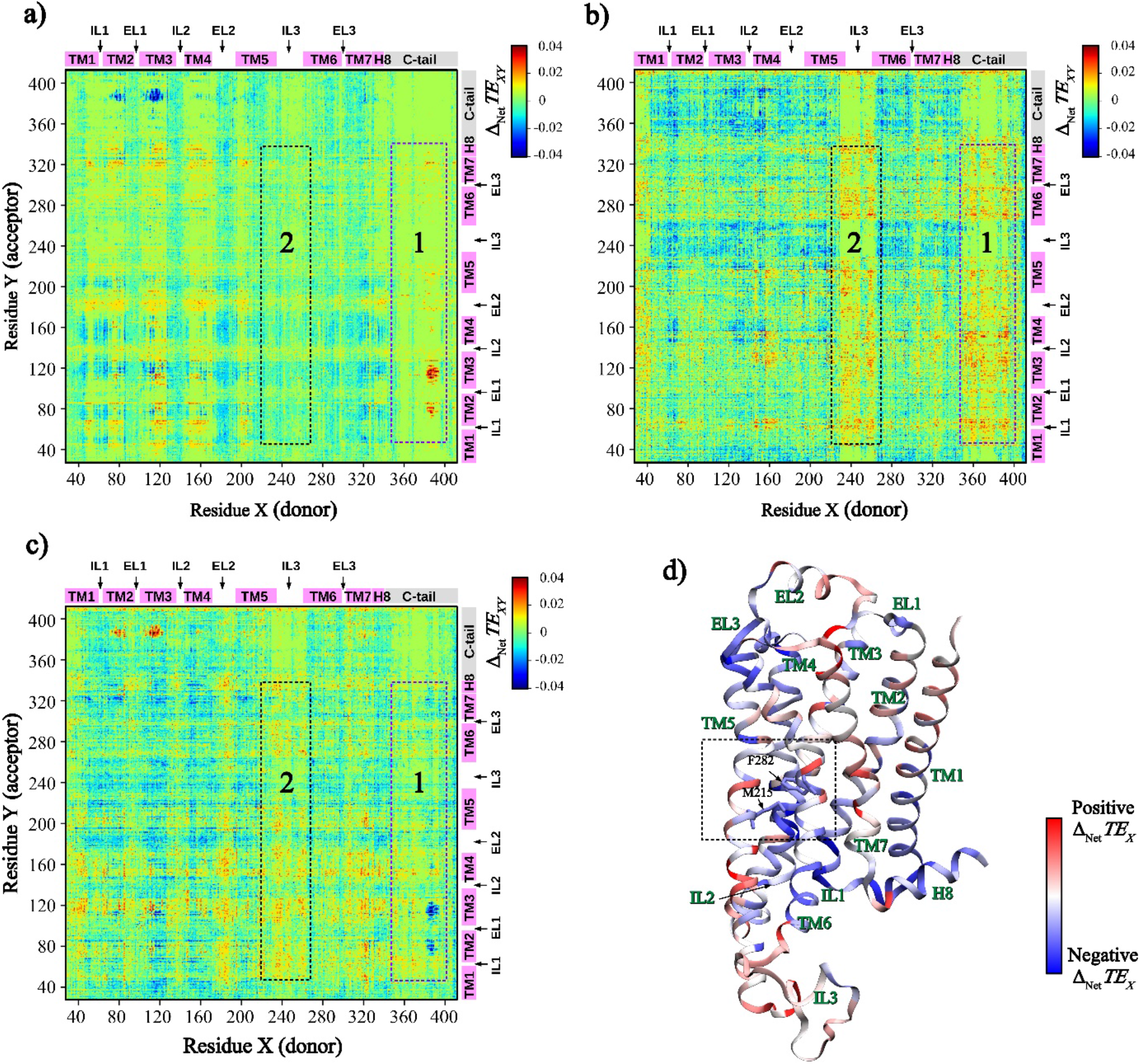
Transfer entropy calculations. Net transfer entropy between residues pairs in **a)** B2AR and **b)** B2ARP. **c)** Change in net transfer entropy upon phosphorylation. **d)** Change in net transfer entropy from each TM residue to all the C-tail residues is shown as a heatmap on the β_2_AR structure.

Further, we observed a significant entropy transfer from the intracellular ends of both TM5, TM6 and their connecting loop IL3 to the rest of the TM domain upon phosphorylation (Figure 4b, c, zone 2). The intracellular half of TM7 as well as H8 acted as entropy donor to various region of TM domain, including residues in TM3, TM4, and intracellular halves of TM1, TM2 and TM5. In addition, EL2 is found to drive correlated motions of different TM helices, mostly, residues in their extracellular halves (Figure 4c). On the other hand, the central parts of TM2, TM3, TM4 and TM5 responded to the correlated motions of various other structural elements of TM regions in the receptor.

Next, we examined the residue level correlated motions in different regions of the receptor and shown in Figure 5. As discussed earlier, the C-tail residues mostly drive the fluctuations of TM residues upon phosphorylation (as shown in Figure 4d). Figure 5a shows a few selected residue pairs with significant *Δ*_Net_*TE*_*XY*_ values, involved in communication from C-tail to the central parts of different TM helices (additional pairs are listed in Table S5). Several residues from TM helices, viz., A46^1.45^, M82^2.53^, T123^3.42^, T164^4.56^, V218^5.57^, and C285^6.47^, accepted entropy from the C-tail. Interestingly, the fluctuations of key residues, M215^5.54^, M279^6.41^, and F282^6.44^ (associated with the structural rearrangements of TM helices in B2ARP)^22^ are found to be driven by the phosphorylated C-tail (Figure 5a). In effect, our results suggest that the allosteric communication from C-tail bring about the changes in conformation in the mid-transmembrane region of β_2_AR upon phosphorylation.

**Figure 5.**
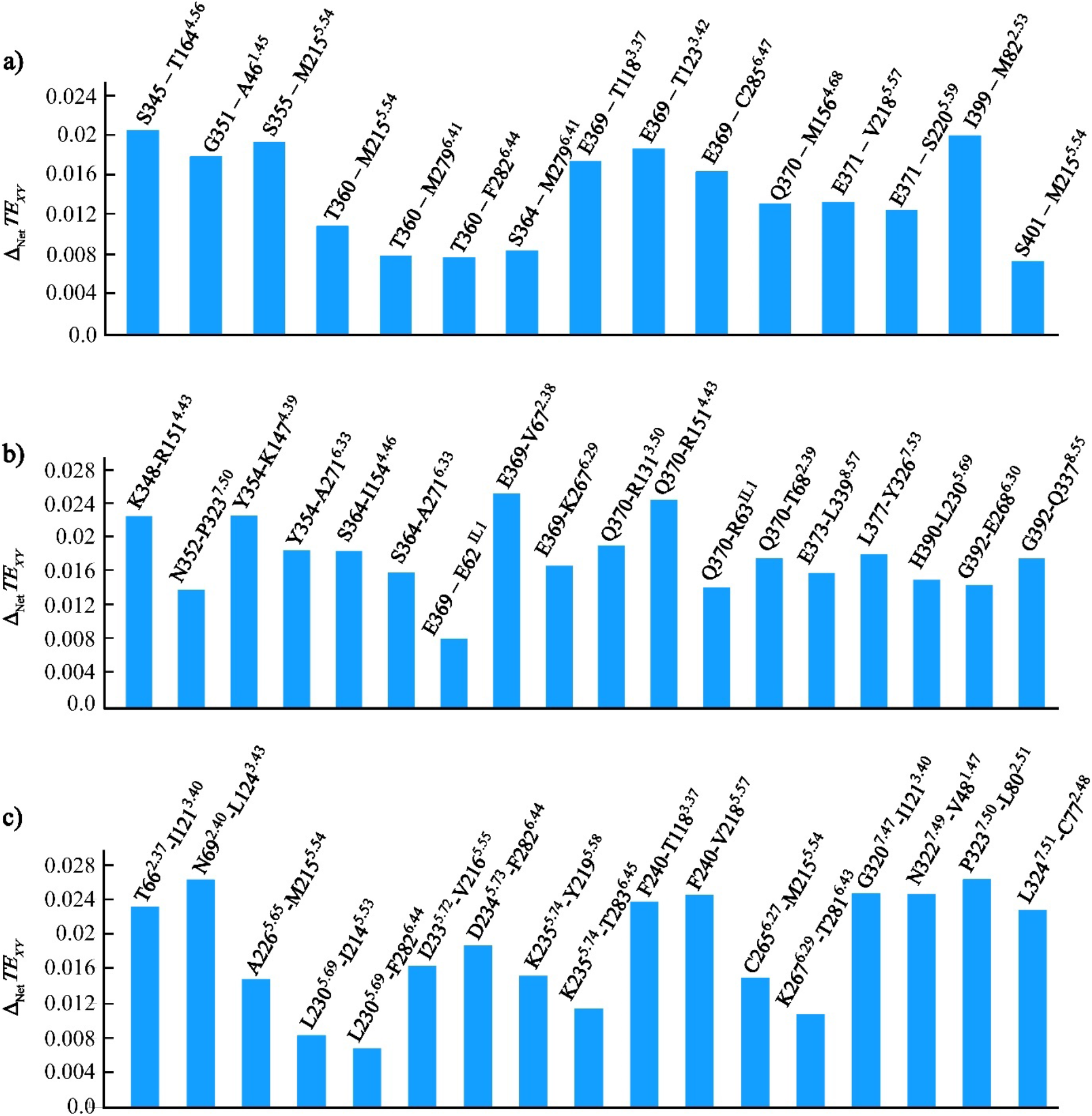
Change in net transfer entropy of selected residue pairs from different regions of β_2_AR upon phosphorylation: **a)** between C-tail and the mid-transmembrane region, **b)** between C-tail and the cytosolic end of the transmembrane domain, and **c)** between cytosolic end of the transmembrane domain and the mid-transmembrane region.

Further, we looked into *Δ*_Net_*TE*_*XY*_ from C-tail to the cytosolic surface of the TM domain (Figure 5b, Table S5). We found that C-tail residues also drive the correlated motions of a number of residues in TM helices and intracellular loops. In fact, several residues from IL1, TM2, TM3, TM4, TM6, and H8 responded strongly to C-tail residues. Particularly, the positively charged residues such as R63^IL1^, R131^3.50^, K147^4.39^, R151^4.43^, and K267^6.29^ are observed to be notable entropy acceptors. In addition, we also found that the TM5 residue L230^5.69^ and the TM7 residues P323^7.50^ and Y326^7.53^ are acting as entropy sinks at the cytosolic surface (Figure 5b, Table S5).

The correlated motions at the cytosolic surface caused by the phosphorylated C-tail, in turn, influenced the residue fluctuations in the mid-transmembrane region of the receptor (Figure 5c, Table S5). Residues from the cytosolic surface including N69^2.40^, R131^3.50^, L230^5.69^, D234^5.73^, F240^IL3^, C265^6.27^, G320^7.47^, and P323^7.50^ are the major entropy donors. Moreover, the positively charged residues, K235^5.74^ and K267^6.29^, also acted as entropy sources of the correlated motions. On the other hand, residues L80^2.51^, T118^3.37^, I121^3.40^, M215^5.54^, V218^5.57^, Y219^5.58^, S220^5.59^, and F282^6.44^ are the significant sinks of entropy (Figure 5c, Table S5), indicating the transfer of information from the cytosolic surface to the mid-transmembrane region in B2ARP. Taken together, our findings therefore suggest that the residues at the cytosolic surface relayed the allosteric signals from the phosphorylated C-tail to the central part of the TM domain of the receptor.

### Redistribution of nonbonding interaction energy network upon phosphorylation

In proteins, rearrangement of inter-residue non-bonding interactions at a region distal to the perturbed site indicates the presence of allosteric communications.^11, 62-65^ Introducing perturbations in terms of phosphorylations at the C-terminus are expected to redistribute the interaction energy network in the receptor.

We calculated the change in average pairwise interaction energies Δ⟨*E*_*XY*_⟩ (eq 12) for all residue pairs in the TM domain from the two conventional MD simulation trajectories (a cumulative length of 1*μ*s) that were started from the minima m_1_ and m_2_ (Figure 1a, b). Δ⟨*E*_*XY*_⟩ values so obtained are divided into seven energy intervals based on their distribution (Figure 6a). Several stabilizing and destabilizing interactions are observed in IL3 and the intracellular ends of TM5 and TM6 (Figure 6a, b, Table S6); in particular, the residue pair E237^IL3^ - K267^6.29^ established the most favourable interaction, whereas, the pair D234^5.73^ - K267^6.29^ is found to be the most unfavourable interaction upon phosphorylation (Figure 6b). Additionally, residue pairs including E225^5.64^ – K232^5.71^, R239^IL3^ – K267^6.29^, and R260^IL3^ – E268^6.30^ show significant stabilization, while the pairs R239^IL3^ – D234^5.73^ and R259^IL3^ – D251^IL3^ are significantly destabilized. Interestingly, the intracellular end of TM5 formed stable interaction with IL2 (K140^IL2^ – D234^5.73^), whereas interactions of TM6 with TM3 and IL2 are destabilized (R131^3.50^ – K270^6.32^, R131^3.50^ – K273^6.35^, and K140^IL2^ – K267^6.29^) upon phosphorylation (Figure 6a, c, Table S6). Furthermore, in the intracellular end of the receptor, H8 residues rearranged their interactions with residues from IL1, TM3, and TM6 (Figure 6a, d, Table S6). It should be noted that the above rearrangements of the interaction energy network are caused due to the conformational variations observed in the cytoplasmic end of the phosphorylated receptor (Figure S5).

**Figure 6.**
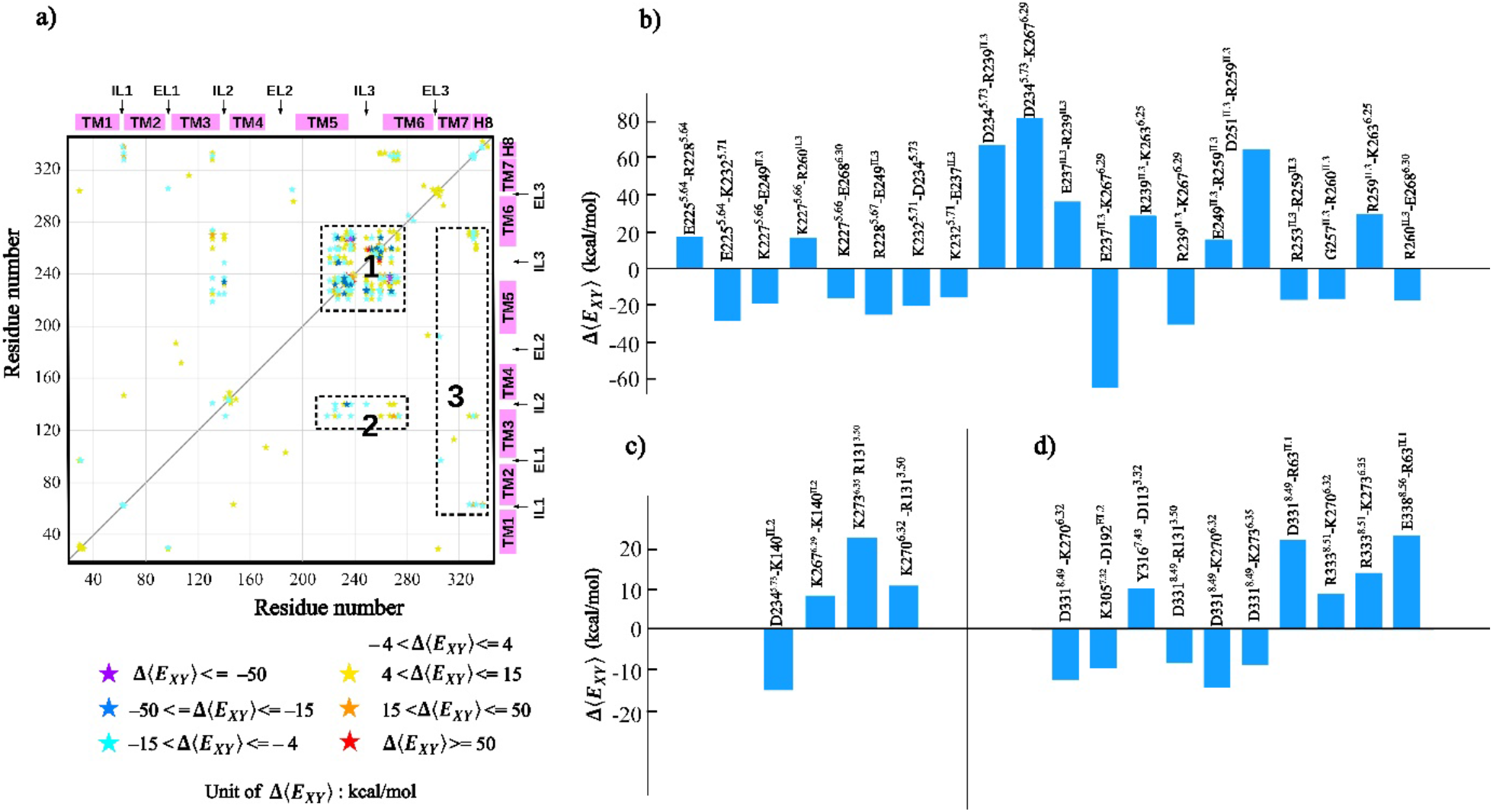
Change in average inter-residue interaction energies upon phosphorylation. **a**) The energy values are divided into seven intervals based on their distribution. Stabilizing interactions or negative energy values are marked in cyan to violet, and destabilizing interactions or positive energy values are marked in yellow to red. **b-d**) Selected significant high energy residue pairs (|Δ⟨*E*_*XY*_⟩| > 8 kcal/mol) from the important zones (1-3) are represented as bar plots.

We further observed that interactions of TM7 with EL1 and EL2 are found to be stabilized and interaction between the ligand-binding residues D113^3.32^ and Y316^7.43^ is destabilized in the extracellular end (Figure 6a, d, Table S6). In fact, such destabilization commensurate with the expansion of the ligand-binding orthosteric pocket observed in B2ARP (Figure S6). Selected residue pairs with significant energy changes shown in Figure 6b, d are also depicted on a representative structure in Figure 7.

**Figure 7.**
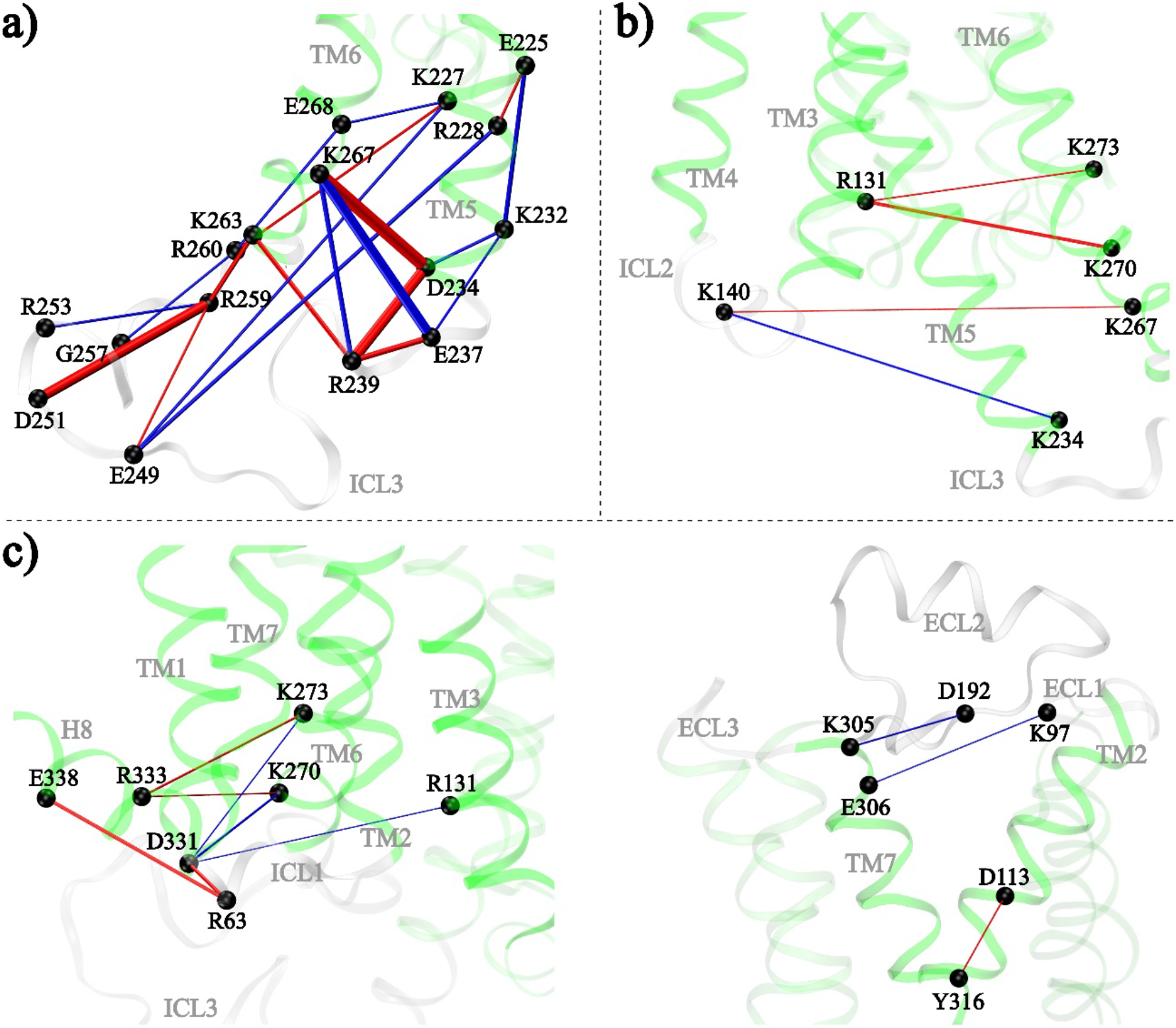
Interaction energy networks within the TM domain. **a**-**c)** Selected significant high energy residue pairs from different zones as shown in Figure 6b-d. Interactions that become favorable upon phosphorylation are shown as blue lines, and those that become unfavorable are shown as red lines. The thickness of lines represents the magnitude of Δ⟨*E*_*XY*_⟩.

It is to be noted that the contribution of the van der Waals interaction energies to the total nonbonding energies are found to be mostly negligible (approximately in the range of -3 to 3 kcal/mol, see Figure S7). However, for residue pairs having comparable electrostatic energies in B2AR and B2ARP systems, change in average pairwise van der Waals energies, 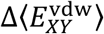, could play a vital role in the redistribution of energy network upon phosphorylation. In particular, residue pairs involving nonpolar and uncharged residues show mostly higher values of 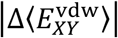 (see Table S7).

We specifically examined the interaction pairs consisting of residues in the mid-transmembrane region, M215^5.54^, F282^6.44^, Y219^5.58^, and M279^6.41^, that NMR data suggested to be allosterically involved in the structural rearrangements of the TM domain.^22^ Figure 8 shows 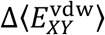 values for M215^5.54^ and Y219^5.58^ with all the other residues. Clearly, the change in interaction energies of the residue pairs M215^5.54^ - F282^6.44^ and M215^5.54^ - M279^6.41^ are positive, indicating the weakening of their interactions upon phosphorylation (Figure 8a). In fact, the destabilizing interaction between M215^5.54^ and F282^6.44^ correlates well with their increased separation observed in B2ARP (Figure 1). Notably, other residue pairs including Y219^5.58^ - M279^6.41^ and Y219^5.58^ - F282^6.44^ also showed positive 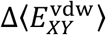, suggesting increased destabilizing interactions in the mid-transmembrane region of the receptor upon phosphorylation (Figure 8b). Such destabilizations demonstrate the allosteric communication from the phosphorylated C-tail to the mid-transmembrane region of the receptor.

**Figure 8.**
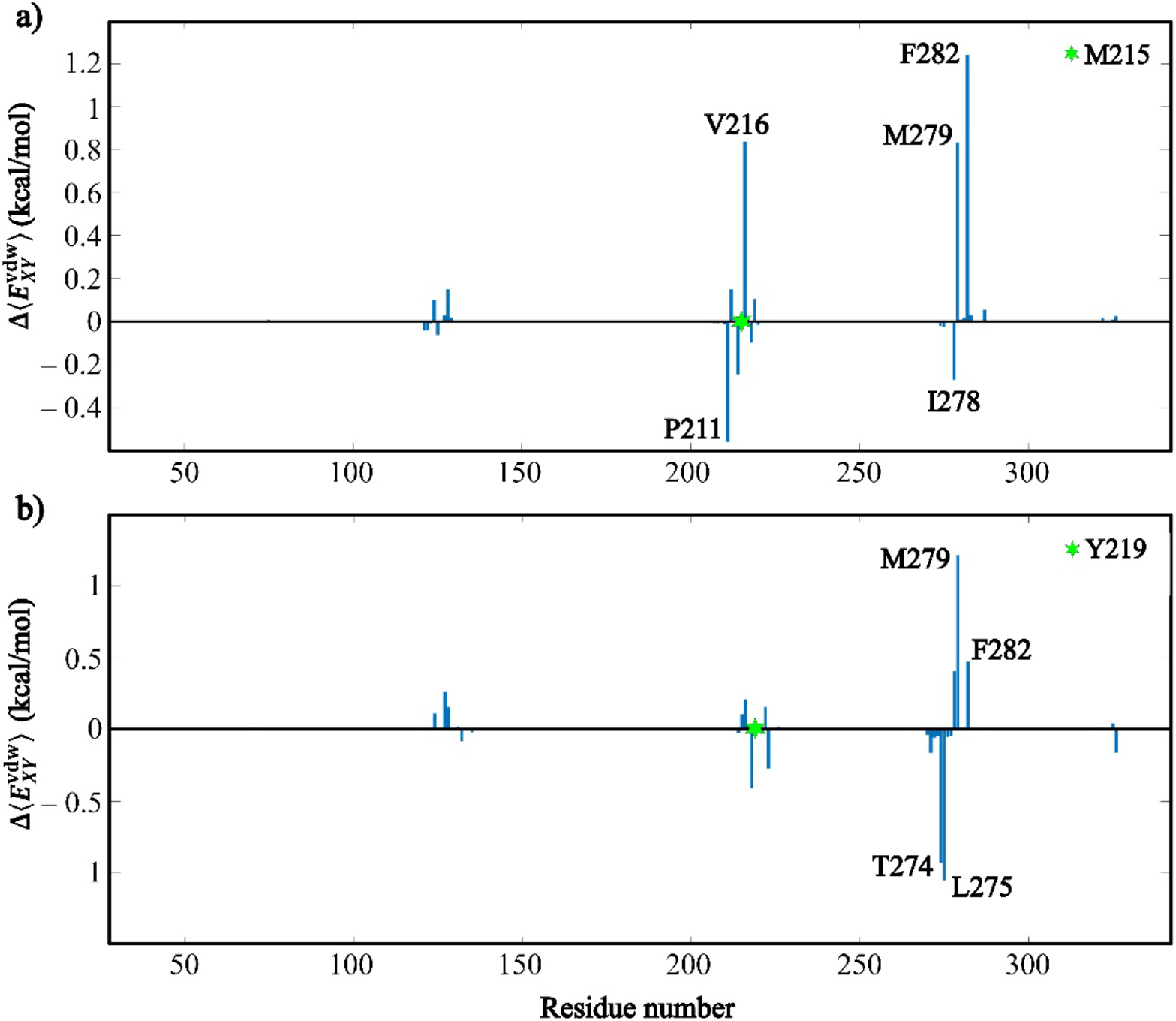
Change in average van der Waals interaction energy 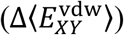 of **a)** M215^5.54^ and **b)** Y219^5.58^ with all the other residues in the TM domain of the receptor.

## Conclusion

In this study, we performed large-scale all-atom conventional and enhanced sampling GaMD simulations to analyze the allosteric effect of phosphorylations at the C-terminus of β_2_AR by GRK2. Compared to the unphosphorylated control, the free energy landscape of the phosphorylated receptor sampled an additional stable conformational state characterized by an inward shift of TM6 at the cytosolic end and separation of the TM5 residue M215^5.54^ from the TM6 residue F282^6.44^ in the mid-transmembrane region. The structural rearrangements at the cytosolic end of the phosphorylated receptor caused a reduction in size of its transducer binding cavity. The efficient binding of a β-arrestin is known to require a narrower cleft at the cytosolic surface in comparison to a G-protein, and therefore, the reduction in cavity volume suggest that the conformational state sampled by the receptor upon phosphorylation is likely to favor β-arrestin over G-protein. The adherence of the phosphorylated C-terminus with a cluster of positively charged residues at the cytosolic surface aids the allosteric communication to the transmembrane domain of the receptor. Our results suggested that the C-terminus upon phosphorylation in general drives the correlated motions of residues in the transmembrane domain and the cytosolic surface residues relayed the allosteric signals. The rewiring of interaction energy networks in the transmembrane domain further demonstrated the long-range communications from the phosphorylated C-terminus. In summary, the residue level insights presented here can aid in rational design of drugs with lesser side effects and superior efficacy.

## Associated Content

### Supporting Information

Figures S1−S7 and Tables S1-S7, providing additional details concerning the results obtained in this study. This material is available free of charge via the Internet at http://pubs.acs.org.

### Notes

The authors declare no competing financial interest.

## Acknowledgement

We acknowledge the high-performance computing facility of IISER Bhopal. M.K.M. is supported by the research fellowships provided by the Council of Scientific & Industrial Research (CSIR), India. A. D. is supported by the research fellowships provided by IISER Bhopal. R. K. M. gratefully acknowledges the financial support provided by Science and Engineering Research Board (SERB), Department of Science and Technology, India (File No. EMR/2016/006815).

---

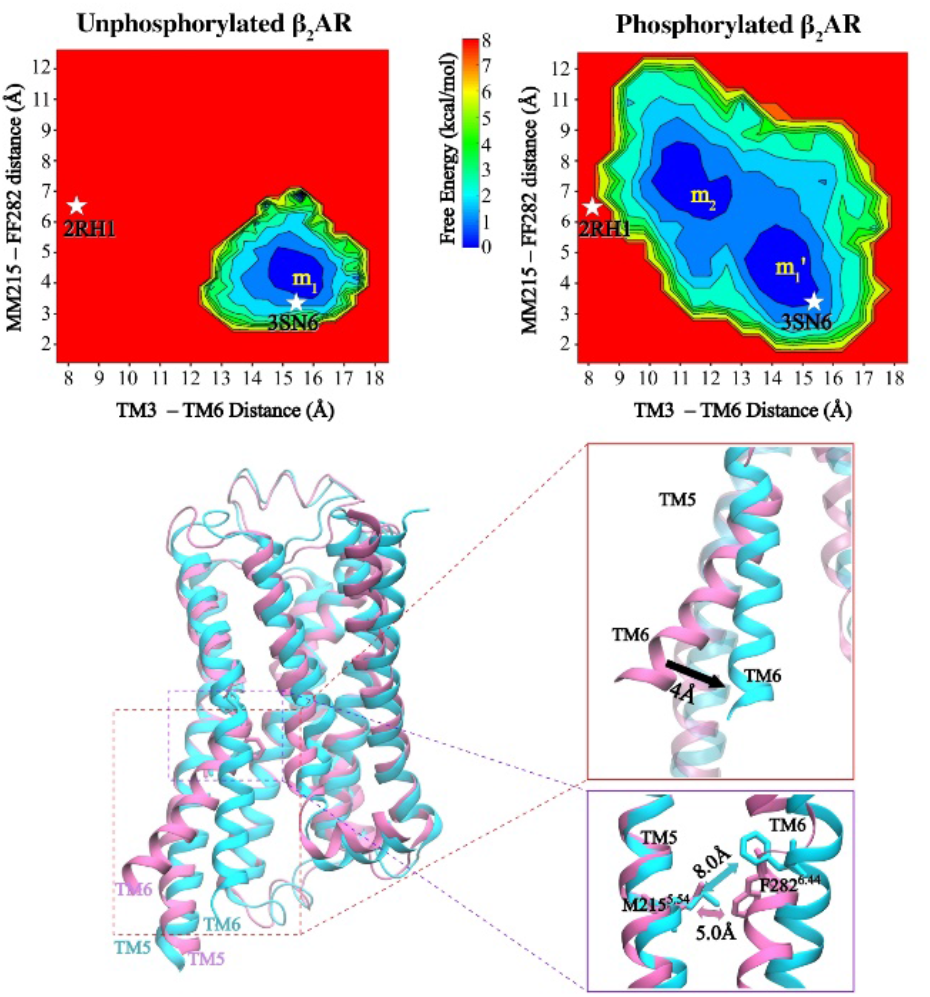

For Table of Contents Only

## References

1. Hilger, D.; Masureel, M.; Kobilka, B. K., Structure and dynamics of GPCR signaling complexes. Nature Structural & Molecular Biology 2018, 25 (1), 4–12.

2. Wacker, D.; Stevens, R. C.; Roth, B. L., How ligands illuminate GPCR molecular pharmacology. Cell 2017, 170 (3), 414–427.

3. Plouffe, B.; Thomsen, A. R.; Irannejad, R., Emerging role of compartmentalized G protein-coupled receptor signaling in the cardiovascular field. ACS Pharmacology & Translational Science 2020, 3 (2), 221–236.

4. Huang, Y.; Todd, N.; Thathiah, A., The role of GPCRs in neurodegenerative diseases: avenues for therapeutic intervention. Current Opinion in Pharmacology 2017, 32, 96–110.

5. Cabral-Marques, O.; Marques, A.; Giil, L. M.; De Vito, R.; Rademacher, J.; Günther, J.; Lange, T.; Humrich, J. Y.; Klapa, S.; Schinke, S., GPCR-specific autoantibody signatures are associated with physiological and pathological immune homeostasis. Nature Communications 2018, 9 (1), 1–14.

6. Gad, A. A.; Balenga, N., The emerging role of adhesion GPCRs in cancer. ACS Pharmacology & Translational Science 2020, 3 (1), 29–42.

7. Yao, X. J.; Ruiz, G. V.; Whorton, M. R.; Rasmussen, S. G.; DeVree, B. T.; Deupi, X.; Sunahara, R. K.; Kobilka, B., The effect of ligand efficacy on the formation and stability of a GPCR-G protein complex. Proceedings of the National Academy of Sciences 2009, 106 (23), 9501–9506.

8. Kenakin, T., A holistic view of GPCR signaling. Nature Biotechnology 2010, 28 (9), 928–929.

9. Weis, W. I.; Kobilka, B. K., The molecular basis of G protein–coupled receptor activation. Annual Review of Biochemistry 2018, 87, 897–919.

10. Du, Y.; Duc, N. M.; Rasmussen, S. G.; Hilger, D.; Kubiak, X.; Wang, L.; Bohon, J.; Kim, H. R.; Wegrecki, M.; Asuru, A., Assembly of a GPCR-G protein complex. Cell 2019, 177 (5), 1232-1242. e11.

11. Singh, R.; Ahalawat, N.; Murarka, R. K., Activation of corticotropin-releasing factor 1 receptor: insights from molecular dynamics simulations. The Journal of Physical Chemistry B 2015, 119 (7), 2806–2817.

12. Lefkowitz, R. J.; Shenoy, S. K., Transduction of receptor signals by ß-arrestins. Science 2005, 308 (5721), 512–517.

13. Gurevich, V. V.; Gurevich, E. V., The structural basis of arrestin-mediated regulation of G-protein-coupled receptors. Pharmacology & Therapeutics 2006, 110 (3), 465–502.

14. Nobles, K. N.; Xiao, K.; Ahn, S.; Shukla, A. K.; Lam, C. M.; Rajagopal, S.; Strachan, R. T.; Huang, T.-Y.; Bressler, E. A.; Hara, M. R., Distinct phosphorylation sites on the β2-adrenergic receptor establish a barcode that encodes differential functions of β-arrestin. Science Signaling 2011, 4 (185), ra51–ra51.

15. Gurevich, E. V.; Gurevich, V. V., Arrestins: ubiquitous regulators of cellular signaling pathways. Genome Biology 2006, 7 (9), 1–10.

16. Gross, O. P.; Burns, M. E., Control of rhodopsin’s active lifetime by arrestin-1 expression in mammalian rods. Journal of Neuroscience 2010, 30 (9), 3450–3457.

17. Gurevich, V. V.; Gurevich, E. V., Biased GPCR signaling: Possible mechanisms and inherent limitations. Pharmacology & Therapeutics 2020, 211, 107540.

18. Huang, W.; Masureel, M.; Qu, Q.; Janetzko, J.; Inoue, A.; Kato, H. E.; Robertson, M. J.; Nguyen, K. C.; Glenn, J. S.; Skiniotis, G., Structure of the neurotensin receptor 1 in complex with β-arrestin 1. Nature 2020, 579 (7798), 303–308.

19. Staus, D. P.; Hu, H.; Robertson, M. J.; Kleinhenz, A. L.; Wingler, L. M.; Capel, W. D.; Latorraca, N. R.; Lefkowitz, R. J.; Skiniotis, G., Structure of the M2 muscarinic receptor–β-arrestin complex in a lipid nanodisc. Nature 2020, 579 (7798), 297–302.

20. Lee, Y.; Warne, T.; Nehmé, R.; Pandey, S.; Dwivedi-Agnihotri, H.; Chaturvedi, M.; Edwards, P. C.; García-Nafría, J.; Leslie, A. G.; Shukla, A. K., Molecular basis of β-arrestin coupling to formoterol-bound β 1-adrenoceptor. Nature 2020, 583 (7818), 862–866.

21. Ballesteros, J. A.; Weinstein, H., [19] Integrated methods for the construction of three-dimensional models and computational probing of structure-function relations in G protein-coupled receptors. In Methods in Neurosciences, Elsevier: 1995; Vol. 25, pp 366–428.

22. Shiraishi, Y.; Natsume, M.; Kofuku, Y.; Imai, S.; Nakata, K.; Mizukoshi, T.; Ueda, T.; Iwaï, H.; Shimada, I., Phosphorylation-induced conformation of β 2-adrenoceptor related to arrestin recruitment revealed by NMR. Nature Communications 2018, 9 (1), 1–10.

23. Miao, Y.; Feher, V. A.; McCammon, J. A., Gaussian accelerated molecular dynamics: Unconstrained enhanced sampling and free energy calculation. Journal of Chemical Theory and Computation 2015, 11 (8), 3584–3595.

24. Wang, J.; Arantes, P. R.; Bhattarai, A.; Hsu, R. V.; Pawnikar, S.; Huang, Y. m. M.; Palermo, G.; Miao, Y., Gaussian accelerated molecular dynamics: Principles and applications. Wiley Interdisciplinary Reviews: Computational Molecular Science 2021, e1521.

25. Rasmussen, S. G.; DeVree, B. T.; Zou, Y.; Kruse, A. C.; Chung, K. Y.; Kobilka, T. S.; Thian, F. S.; Chae, P. S.; Pardon, E.; Calinski, D., Crystal structure of the β 2 adrenergic receptor–Gs protein complex. Nature 2011, 477 (7366), 549–555.

26. Webb, B.; Sali, A., Comparative protein structure modeling using MODELLER. Current Protocols in Bioinformatics 2016, 54 (1), 5.6. 1-5.6. 37.

27. Cherezov, V.; Rosenbaum, D. M.; Hanson, M. A.; Rasmussen, S. G.; Thian, F. S.; Kobilka, T. S.; Choi, H.-J.; Kuhn, P.; Weis, W. I.; Kobilka, B. K., High-resolution crystal structure of an engineered human β2-adrenergic G protein– coupled receptor. Science 2007, 318 (5854), 1258–1265.

28. Drozdetskiy, A.; Cole, C.; Procter, J.; Barton, G. J., JPred4: a protein secondary structure prediction server. Nucleic Acids Research 2015, 43 (W1), W389–W394.

29. McGuffin, L. J.; Bryson, K.; Jones, D. T., The PSIPRED protein structure prediction server. Bioinformatics 2000, 16 (4), 404–405.

30. Venkatakrishnan, A.; Flock, T.; Prado, D. E.; Oates, M. E.; Gough, J.; Babu, M. M., Structured and disordered facets of the GPCR fold. Current Opinion in Structural Biology 2014, 27, 129–137.

31. Dror, R. O.; Arlow, D. H.; Maragakis, P.; Mildorf, T. J.; Pan, A. C.; Xu, H.; Borhani, D. W.; Shaw, D. E., Activation mechanism of the β2-adrenergic receptor. Proceedings of the National Academy of Sciences 2011, 108 (46), 18684–18689.

32. Vanommeslaeghe, K.; Hatcher, E.; Acharya, C.; Kundu, S.; Zhong, S.; Shim, J.; Darian, E.; Guvench, O.; Lopes, P.; Vorobyov, I., CHARMM general force field: A force field for drug-like molecules compatible with the CHARMM all-atom additive biological force fields. Journal of Computational Chemistry 2010, 31 (4), 671–690.

33. Vanommeslaeghe, K.; MacKerell Jr, A. D., Automation of the CHARMM General Force Field (CGenFF) I: bond perception and atom typing. Journal of Chemical Information and Modeling 2012, 52 (12), 3144–3154.

34. Wu, E. L.; Cheng, X.; Jo, S.; Rui, H.; Song, K. C.; Dávila-Contreras, E. M.; Qi, Y.; Lee, J.; Monje-Galvan, V.; Venable, R. M., CHARMM-GUI membrane builder toward realistic biological membrane simulations. Wiley Online Library: 2014.

35. Jo, S.; Kim, T.; Im, W., Automated builder and database of protein/membrane complexes for molecular dynamics simulations. PloS One 2007, 2 (9), e880.

36. Huang, J.; Rauscher, S.; Nawrocki, G.; Ran, T.; Feig, M.; De Groot, B. L.; Grubmüller, H.; MacKerell, A. D., CHARMM36m: an improved force field for folded and intrinsically disordered proteins. Nature Methods 2017, 14 (1), 71–73.

37. Huang, J.; MacKerell Jr, A. D., CHARMM36 all-atom additive protein force field: Validation based on comparison to NMR data. Journal of Computational Chemistry 2013, 34 (25), 2135–2145.

38. Boonstra, S.; Onck, P. R.; van der Giessen, E., CHARMM TIP3P water model suppresses peptide folding by solvating the unfolded state. The Journal of Physical Chemistry B 2016, 120 (15), 3692–3698.

39. Jo, S.; Cheng, X.; Islam, S. M.; Huang, L.; Rui, H.; Zhu, A.; Lee, H. S.; Qi, Y.; Han, W.; Vanommeslaeghe, K., CHARMM-GUI PDB manipulator for advanced modeling and simulations of proteins containing nonstandard residues. Advances in Protein Chemistry and Structural Biology 2014, 96, 235–265.

40. Liu, W.; Schmidt, B.; Voss, G.; Müller-Wittig, W., Accelerating molecular dynamics simulations using Graphics Processing Units with CUDA. Computer Physics Communications 2008, 179 (9), 634–641.

41. Yang, Y. I.; Shao, Q.; Zhang, J.; Yang, L.; Gao, Y. Q., Enhanced sampling in molecular dynamics. The Journal of Chemical Physics 2019, 151 (7), 070902.

42. Saleh, N.; Saladino, G.; Gervasio, F. L.; Clark, T., Investigating allosteric effects on the functional dynamics of β2-adrenergic ternary complexes with enhanced-sampling simulations. Chemical Science 2017, 8 (5), 4019–4026.

43. Ahalawat, N.; Arora, S.; Murarka, R. K., Structural Ensemble of CD4 Cytoplasmic Tail (402–419) Reveals a Nearly Flat Free-Energy Landscape with Local α-Helical Order in Aqueous Solution. The Journal of Physical Chemistry B 2015, 119 (34), 11229–11242.

44. Miao, Y.; McCammon, J. A., Graded activation and free energy landscapes of a muscarinic G-protein–coupled receptor. Proceedings of the National Academy of Sciences 2016, 113 (43), 12162–12167.

45. Miao, Y.; Bhattarai, A.; Nguyen, A. T.; Christopoulos, A.; May, L. T., Structural basis for binding of allosteric drug leads in the adenosine A 1 receptor. Scientific Reports 2018, 8 (1), 1–13.

46. Miao, Y.; McCammon, J. A., Mechanism of the G-protein mimetic nanobody binding to a muscarinic G-protein-coupled receptor. Proceedings of the National Academy of Sciences 2018, 115 (12), 3036–3041.

47. Miao, Y.; McCammon, J. A., Gaussian accelerated molecular dynamics: Theory, implementation, and applications. In Annual Reports in Computational Chemistry, Elsevier: 2017; Vol. 13, pp 231–278.

48. Schreiber, T., Measuring information transfer. Physical Review Letters 2000, 85 (2), 461.

49. Takens, F., In dynamical systems of turbulence. Lecture Notes in Mathematics 1981, 898, 366.

50. Wibral, M.; Pampu, N.; Priesemann, V.; Siebenhühner, F.; Seiwert, H.; Lindner, M.; Lizier, J. T.; Vicente, R., Measuring information-transfer delays. PloS One 2013, 8 (2), e55809.

51. Lindner, M.; Vicente, R.; Priesemann, V.; Wibral, M., TRENTOOL: A Matlab open source toolbox to analyse information flow in time series data with transfer entropy. BMC Neuroscience 2011, 12 (1), 1–22.

52. Kraskov, A.; Stögbauer, H.; Grassberger, P., Estimating mutual information. Physical Review E 2004, 69 (6), 066138.

53. Ragwitz, M.; Kantz, H., Markov models from data by simple nonlinear time series predictors in delay embedding spaces. Physical Review E 2002, 65 (5), 056201.

54. Marschinski, R.; Kantz, H., Analysing the information flow between financial time series. The European Physical Journal B-Condensed Matter and Complex Systems 2002, 30 (2), 275–281.

55. Kamberaj, H.; van der Vaart, A., Extracting the causality of correlated motions from molecular dynamics simulations. Biophysical Journal 2009, 97 (6), 1747–1755.

56. McGibbon, R. T.; Beauchamp, K. A.; Harrigan, M. P.; Klein, C.; Swails, J. M.; Hernández, C. X.; Schwantes, C. R.; Wang, L.-P.; Lane, T. J.; Pande, V. S., MDTraj: a modern open library for the analysis of molecular dynamics trajectories. Biophysical Journal 2015, 109 (8), 1528–1532.

57. Chakir, K.; Xiang, Y.; Yang, D.; Zhang, S.-J.; Cheng, H.; Kobilka, B. K.; Xiao, R.-P., The third intracellular loop and the carboxyl terminus of β2-adrenergic receptor confer spontaneous activity of the receptor. Molecular Pharmacology 2003, 64 (5), 1048–1058.

58. Lu, Z.-L.; Saldanha, J. W.; Hulme, E. C., Seven-transmembrane receptors: crystals clarify. Trends in Pharmacological Sciences 2002, 23 (3), 140–146.

59. Kostenis, E.; Conklin, B. R.; Wess, J., Molecular basis of receptor/G protein coupling selectivity studied by coexpression of wild type and mutant m2 muscarinic receptors with mutant Gαq subunits. Biochemistry 1997, 36 (6), 1487–1495.

60. Kunkel, M. T.; Peralta, E. G., Charged amino acids required for signal transduction by the m3 muscarinic acetylcholine receptor. The EMBO Journal 1993, 12 (10), 3809–3815.

61. Dolinsky, T. J.; Nielsen, J. E.; McCammon, J. A.; Baker, N. A., PDB2PQR: an automated pipeline for the setup of Poisson–Boltzmann electrostatics calculations. Nucleic Acids Research 2004, 32 (Suppl_2), W665–W667.

62. Kumawat, A.; Chakrabarty, S., Hidden electrostatic basis of dynamic allostery in a PDZ domain. Proceedings of the National Academy of Sciences 2017, 114 (29), E5825–E5834.

63. Kumawat, A.; Chakrabarty, S., Protonation-Induced Dynamic Allostery in PDZ Domain: Evidence of Perturbation-Independent Universal Response Network. The Journal of Physical Chemistry Letters 2020, 11 (21), 9026–9031.

64. Vijayabaskar, M.; Vishveshwara, S., Interaction energy based protein structure networks. Biophysical journal 2010, 99 (11), 3704–3715.

65. Liu, J.; Nussinov, R., Energetic redistribution in allostery to execute protein function. Proceedings of the National Academy of Sciences 2017, 114 (29), 7480–7482.

