## Supplementary Material for "Molecular insights into phosphorylation-induced allosteric conformational changes in β_2_-adrenergic receptor"

AUTHOR INFORMATION

**Corresponding Author**

*

### Supplementary Figures

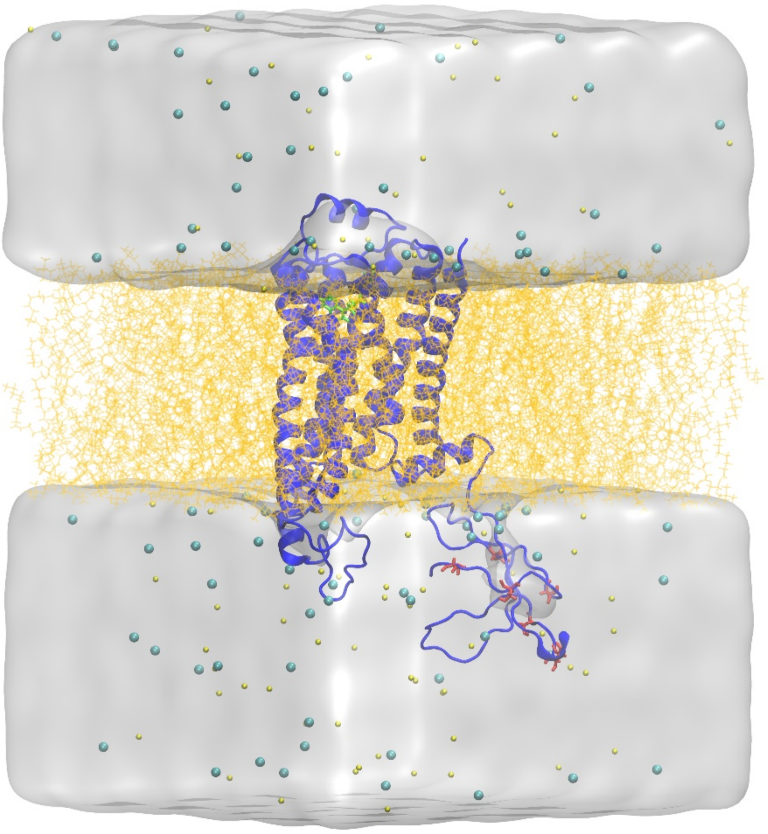

**Figure S1.** Schematic representation of β2AR (blue ribbon) inserted into POPC bilayer (light orange lines) and solvated in water (white transparent surface) neutralized by 150 mM sodium (yellow spheres) and chlorine (cyan spheres). Phosphorylated residues (red licorice) are shown on the C-tail of the receptor.

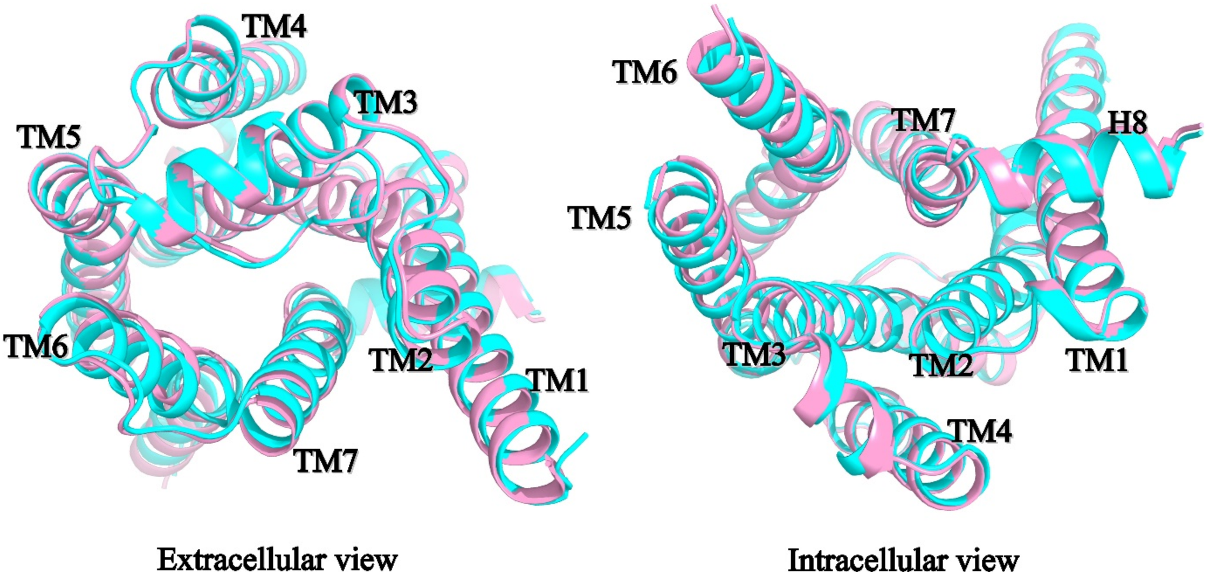


**Figure S2.** Average structures from minima $m_1$ (B2AR; pink) and $m'_1$ (B2ARP; cyan).

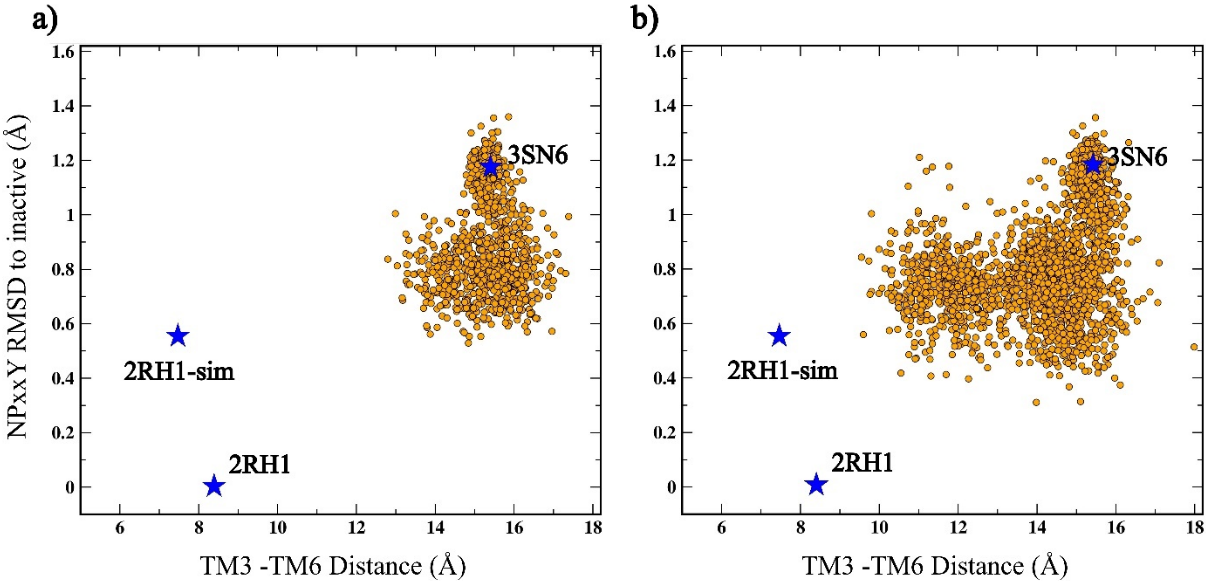


**Figure S3.** GaMD simulation results of **a)** unphosphorylated and **b)** phosphorylated receptors. 2RH1-sim is the mean of CVs of the inactive state cluster observed in the simulation. The scattered plot of CVs was chosen to compare the results with the previous simulation study mentioned in the main text.

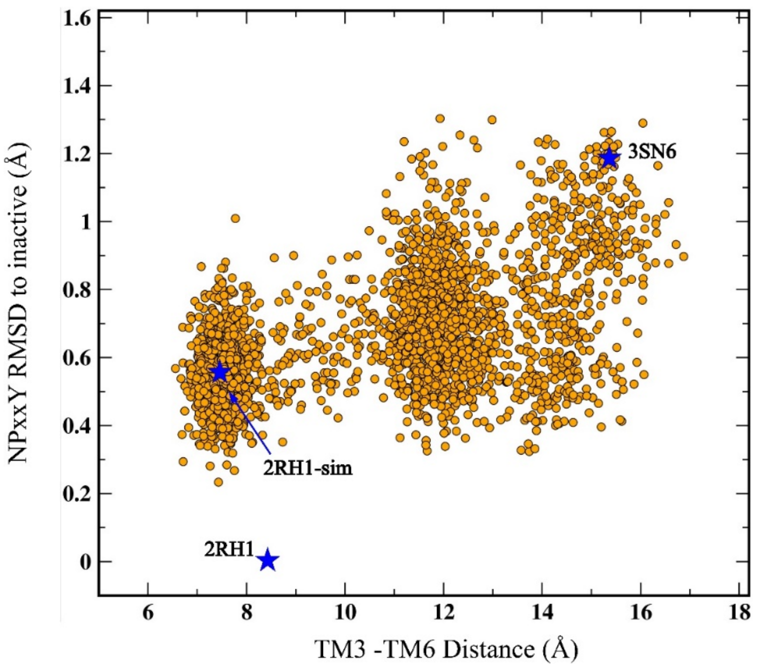


**Figure S4.** GaMD simulation results of truncated β2AR. The system started from the active structure (PDB ID: 3SN6) sampled inactive state conformations (similar to PDB ID 2RH1). 2RH1-sim is the mean of CVs of the inactive state cluster observed in the simulation. The scattered plot of CVs was chosen to compare the results with the previous simulation study mentioned in the main text.

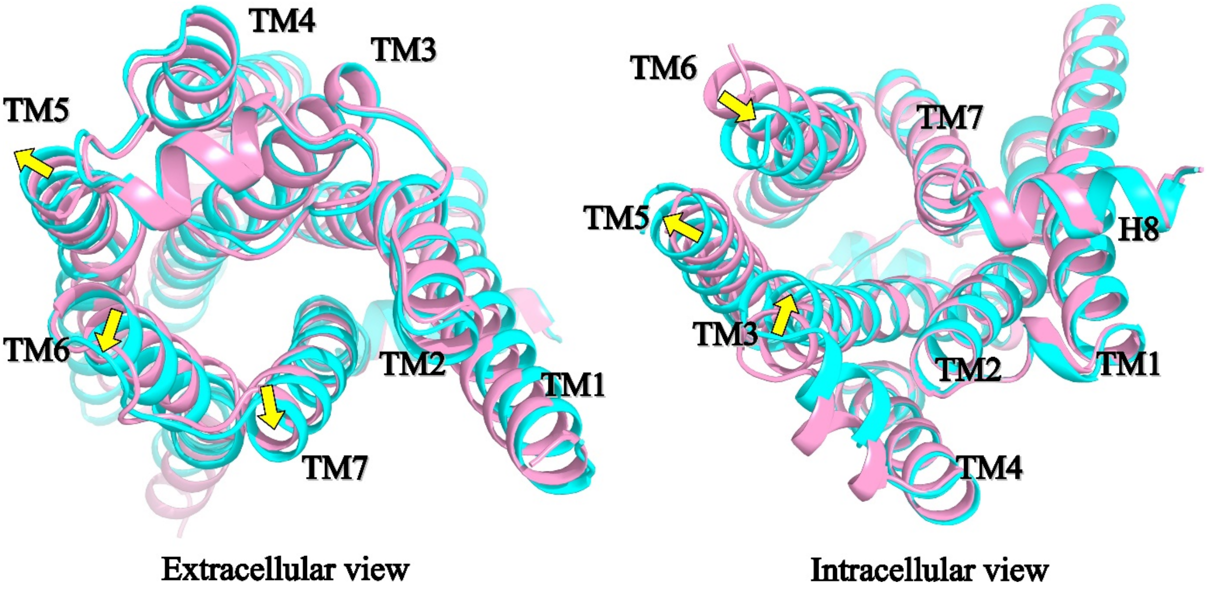


**Figure S5.** Average structures from minima $m_1$ (B2AR; pink) and $m_2$ (B2ARP; cyan). The average structural differences are marked using yellow arrows.

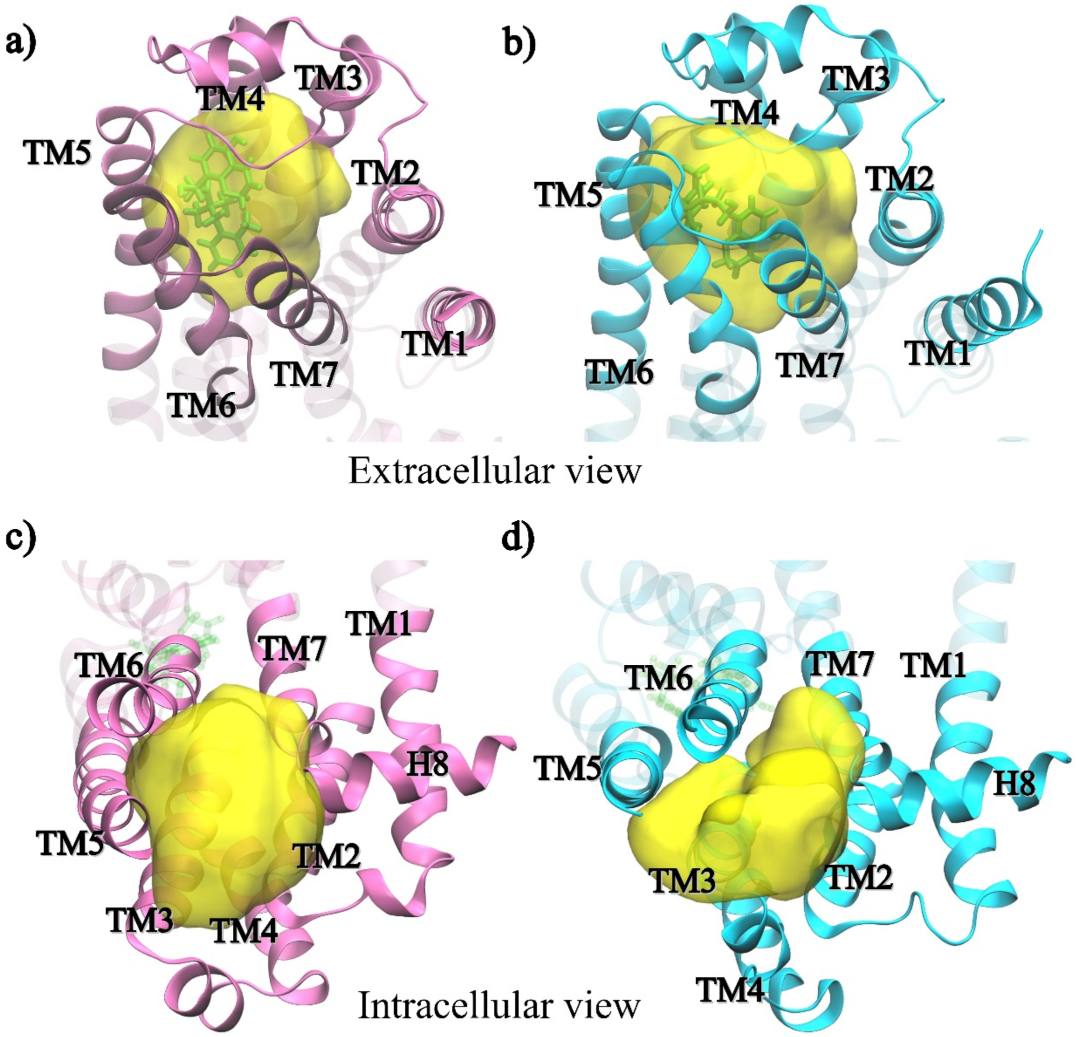


**Figure S6.** Volumes of receptor cavities (yellow transparent surface) of B2AR (pink) and B2ARP (cyan) calculated using POVME 3.0. **a, b)** Extracellular orthosteric pocket of B2AR and B2ARP. **c, d)** Intracellular transducer binding cavity of B2AR and B2ARP.

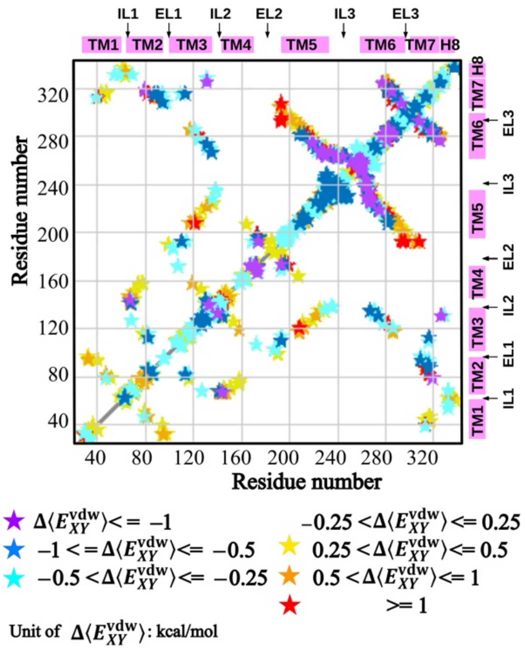


**Figure S7.** Change in average van der Waals ($\Delta<E_{XY}^{vdW}>$) interaction energies of all residue pairs color coded for better understanding.

### Supplementary tables

**Table S1.** GaMD simulation summary of both unphosphorylated and phosphorylated systems. Average and standard deviation of total boost potential ($\Delta V$) is given.

| Simulation ID | Length (ns) | $\Delta V$(kcal/mol) | |
|---|---|---|---|
| | | Avg | σ |
| Unphosphorylated system | | | |
| SIM1 | 2000 | 15.85 | 4.64 |
| SIM2 | 2000 | 15.78 | 4.62 |
| SIM3 | 2000 | 15.82 | 4.63 |
| Phosphorylated system | | | |
| SIM1 | 2000 | 15.87 | 4.65 |
| SIM2 | 2000 | 15.96 | 4.67 |
| SIM3 | 2000 | 15.94 | 4.66 |

**Table S2.** Residue details of various receptor regions and structural elements.

| Receptor region | Residues | Extracellular end | Intracellular end | Centre part |
|---|---|---|---|---|
| TM1 | $Q26^{1.25}$-$F61^{1.60}$ | $Q26^{1.25}$-$V33^{1.32}$ | $V54^{1.53}$-$F61^{1.60}$ | $S41^{1.40}$-$V48^{1.47}$ |
| IL1 | $E62^{IL1}$ - $Q65^{IL1}$ | - | - | - |
| TM2 | $T66^{2.37}$-$M96^{2.67}$ | $F89^{2.60}$-$M96^{2.67}$ | $T66^{2.37}$-$T73^{2.44}$ | $A78^{2.49}$-$A85^{2.56}$ |
| EL1 | $K97^{EL1}$ - $F101^{EL1}$ | - | - | - |
| TM3 | $G102^{3.21}$-$S137^{3.56}$ | $G102^{3.21}$-$W109^{3.28}$ | $D130^{3.49}$-$S137^{3.56}$ | $V117^{3.36}$-$L124^{3.43}$ |
| IL2 | $P138^{IL2}$ - $L145^{IL2}$ | - | - | - |
| TM4 | $T146^{4.38}$-$H172^{4.64}$ | $S165^{4.57}$-$H172^{4.64}$ | $T146^{4.38}$-$I153^{4.45}$ | $M156^{4.48}$-$L163^{4.55}$ |
| EL2 | $W173^{EL2}$ - $T195^{EL2}$ | - | - | - |
| TM5 | $N196^{5.35}$-$K235^{5.74}$ | $N196^{5.35}$-$S203^{5.42}$ | $R228^{5.67}$-$K235^{5.74}$ | $V213^{5.52}$-$S220^{5.59}$ |
| IL3 | $S236^{IL3}$ - $S261^{IL3}$ | - | - | - |
| TM6 | $S262^{6.24}$-$Q299^{6.61}$ | $V292^{6.54}$-$Q299^{6.61}$ | $S262^{6.24}$-$H269^{6.31}$ | $I278^{6.40}$-$C285^{6.47}$ |
| EL3 | $D300^{EL3}$ - $I303^{EL3}$ | - | - | - |
| TM7 | $R304^{7.31}$-$R328^{7.55}$ | $R304^{7.31}$-$L311^{7.38}$ | $F321^{7.48}$-$R328^{7.55}$ | $W313^{7.40}$-$G320^{7.47}$ |
| H8 | $S329^{8.47}$ - $C341^{8.59}$ | - | - | - |
| C-tail | L342 - L413 | - | - | - |

**Table S3.** Unique contacts in B2AR and B2ARP. A contact between two residues is defined when any heavy atom of one residue comes <5Å closer to any atom in another residue for 80% cMD trajectory frames.

| **Regions** | **Residues forming unique contacts** |
|---|---|
| Between TM region and C-tail in B2AR | ($R131^{3.50}$, L413) |
| Between TM region and C-tail in B2ARP | ($F61^{1.60}$, V388), ($R63^{\mathrm{IL1}}$, D386), ($R63^{\mathrm{IL1}}$, V388), ($L64^{\mathrm{IL1}}$, V388), ($R259^{\mathrm{IL3}}$, S407), ($R260^{\mathrm{IL3}}$, N409), ($S261^{\mathrm{IL3}}$, N409), ($S262^{6.24}$, N409), ($K263^{6.25}$, Y350), ($K263^{6.25}$, T360), ($L266^{6.28}$, Y350), ($L266^{6.28}$, N409), ($H269^{6.31}$, N409), ($K270^{6.32}$, S411), ($K273^{6.35}$, D410), ($K273^{6.35}$, S411), ($P330^{8.48}$, L413), ($D331^{8.49}$, V388), ($R333^{8.51}$, L413), ($I334^{8.52}$, F387), ($I334^{8.52}$, V388), ($I334^{8.52}$, L413), ($A335^{8.53}$, V388), ($E338^{8.56}$, R344) |
| Within TM region in B2AR | ($D29^{1.28}$, $W32^{1.31}$), ($D29^{1.28}$, $V33^{1.32}$), ($W32^{1.31}$, $G37^{1.36}$), ($M36^{1.35}$, $L95^{2.66}$), ($M40^{1.39}$, $W313^{7.40}$), ($M40^{1.39}$, $I94^{2.65}$), ($I43^{1.42}$, $W313^{7.40}$), ($I43^{1.42}$, $V317^{7.44}$), ($L45^{1.44}$, $G50^{1.49}$), ($I47^{1.46}$, $Y316^{7.43}$), ($I47^{1.46}$, $M82^{2.53}$), ($V48^{1.47}$, $G83^{2.54}$), ($I58^{1.57}$, $Y70^{2.41}$), ($A59^{1.58}$, $Q65^{\mathrm{IL1}}$), ($Q65^{\mathrm{IL1}}$, $Y70^{2.41}$), ($V67^{2.38}$, $Y141^{\mathrm{IL2}}$), ($V67^{2.38}$, $T146^{4.38}$), ($V67^{2.38}$, $A150^{4.42}$), ($Y70^{2.41}$, $K147^{4.39}$), ($Y70^{2.41}$, $A150^{4.42}$), ($F71^{2.42}$, $D130^{3.49}$), ($F71^{2.42}$, A1504.42), (S742.45, T1233.42), ($S74^{2.45}$, $W158^{4.50}$), ($L75^{2.46}$, $S120^{3.39}$), ($L75^{2.46}$, $I127^{3.46}$), ($A78^{2.49}$, $W158^{4.50}$), ($V87^{2.58}$, $W313^{7.40}$), ($G90^{2.61}$, $W313^{7.40}$), ($A91^{2.62}$, $W313^{7.40}$), ($H93^{2.64}$, $C191^{\mathrm{IL1}}$), ($M96^{2.67}$, $T100^{\mathrm{IL1}}$), ($E107^{3.26}$, $H172^{4.64}$), ($D113^{3.32}$, $Y316^{7.43}$), ($V117^{3.36}$, $F208^{5.47}$), ($T118^{3.37}$, $S165^{4.57}$), ($S120^{3.39}$, $W286^{6.48}$), ($I121^{3.40}$, $F208^{5.47}$), ($I121^{3.40}$, $L212^{5.51}$), ($E122^{3.41}$, $P211^{5.50}$), ($L124^{3.43}$, $F282^{6.44}$), ($L124^{3.43}$, $W286^{6.48}$), ($V126^{3.45}$, $V157^{4.49}$), ($A128^{3.47}$, $V222^{5.61}$), ($F133^{3.52}$, $P138^{\mathrm{IL1}}$), ($I135^{3.54}$, $A226^{5.65}$), ($Y141^{\mathrm{IL1}}$, $L145^{\mathrm{IL1}}$), ($Q142^{\mathrm{IL1}}$, $L145^{\mathrm{IL1}}$), ($F193^{\mathrm{IL1}}$, $N293^{6.55}$), ($F193^{\mathrm{IL1}}$, $H296^{6.58}$), ($F193^{\mathrm{IL1}}$, $V297^{6.59}$), ($F193^{\mathrm{IL1}}$, $Y308^{7.35}$), ($Q197^{5.36}$, $V297^{6.59}$), ($A200^{5.39}$, $V297^{6.59}$), ($A202^{5.41}$, $S207^{5.46}$), ($S204^{5.43}$, $I294^{6.56}$), ($F208^{5.47}$, $W286^{6.48}$), ($Y209^{5.48}$, $F290^{6.52}$), ($M215^{5.54}$, $F282^{6.44}$), ($V216^{5.55}$, $M279^{6.41}$), ($Y219^{5.58}$, $M279^{6.41}$), ($F223^{5.62}$, $E268^{6.30}$), ($F223^{5.62}$, $L272^{6.34}$), ($F223^{5.62}$, $L275^{6.37}$), ($A226^{5.65}$, $L230^{5.69}$), ($K227^{5.66}$, $L230^{5.69}$), ($K227^{5.66}$, $Q231^{5.70}$), ($Q229^{5.68}$, $I233^{5.72}$), ($L230^{5.69}$, $I233^{5.72}$), ($L230^{5.69}$, $D234^{5.73}$), ($L230^{5.69}$, $K267^{6.29}$), ($Q231^{5.70}$, $D234^{5.73}$), ($Q231^{5.70}$, $K235^{5.74}$), ($K232^{5.71}$, $K235^{5.74}$), ($K232^{5.71}$, $S236^{\mathrm{IL1}}$), ($I233^{5.72}$, $S236^{\mathrm{IL1}}$), ($I233^{5.72}$, $E237^{\mathrm{IL1}}$), ($D234^{5.73}$, $E237^{\mathrm{IL1}}$), ($D234^{5.73}$, $G238^{\mathrm{IL1}}$), ($D234^{5.73}$, $R239^{\mathrm{IL1}}$), ($D234^{5.73}$, $F240^{\mathrm{IL1}}$), ($D234^{5.73}$, $K267^{6.29}$), ($K235^{5.74}$, |

| | |
|---|---|
| | G238$^{IL1}$), (G276$^{6.38}$, T281$^{6.43}$), (T281$^{6.43}$, F321$^{7.48}$), (T281$^{6.43}$, N322$^{7.49}$), (F282$^{6.44}$, N318$^{7.45}$), (F282$^{6.44}$, Y326$^{7.53}$), (T283$^{6.45}$, W286$^{6.48}$), (L284$^{6.46}$, N318$^{7.45}$), (C285$^{6.47}$, L311$^{7.38}$), (C285$^{6.47}$, G315$^{7.42}$), (W286$^{6.48}$, G315$^{7.42}$), (W286$^{6.48}$, S319$^{7.46}$), (F289$^{6.51}$, Y308$^{7.35}$), (V292$^{6.54}$, Y308$^{7.35}$), (V295$^{6.57}$, I303$^{IL1}$), (H296$^{6.58}$, D300$^{IL1}$), (H296$^{6.58}$, Y308$^{7.35}$) |
| Within TM region in B2ARP | (N103$^{3.22}$, C184), (N103$^{3.22}$, A186), (N103$^{3.22}$, N187), (N103$^{3.22}$, C190), (W109$^{3.28}$, Y316$^{7.43}$), (V114$^{3.33}$, Y199$^{5.38}$), (S120$^{3.39}$, N322$^{7.49}$), (Y132$^{3.51}$, E225$^{5.64}$), (F133$^{3.52}$, K140), (F133$^{3.52}$, L144), (T136$^{3.55}$, E225$^{5.64}$), (W173, N196$^{5.35}$), (R175, T195), (A176, F194), (D192, K305$^{7.32}$), (F208$^{5.47}$, F282$^{6.44}$), (L212$^{5.51}$, F282$^{6.44}$), (M215$^{5.54}$, S220$^{5.59}$), (Y219$^{5.58}$, I278$^{6.40}$), (V222$^{5.61}$, A271$^{6.33}$), (A226$^{5.65}$, K267$^{6.29}$), (A226$^{5.65}$, E268$^{6.30}$), (A226$^{5.65}$, A271$^{6.33}$), (E237, K267$^{6.29}$), (H256, R260), (G257, R260), (R260, H269$^{6.31}$), (S262$^{6.24}$, L266$^{6.28}$), (A271$^{6.33}$, G276$^{6.38}$), (I277$^{6.39}$, I325$^{7.52}$), (I278$^{6.40}$, Y326$^{7.53}$), (T281$^{6.43}$, C285$^{6.47}$), (T281$^{6.43}$, W286$^{6.48}$), (F282$^{6.44}$, C285$^{6.47}$), (F282$^{6.44}$, P288$^{6.50}$), (F289$^{6.51}$, G315$^{7.42}$), (H296$^{6.58}$, L302), (C327$^{7.54}$, F336$^{8.54}$), (L53$^{1.52}$, L339$^{8.57}$), (V54$^{1.53}$, F332$^{8.50}$), (V54$^{1.53}$, T73$^{2.44}$), (A57$^{1.56}$, A335$^{8.53}$), (A57$^{1.56}$, L339$^{8.57}$), (K60$^{1.59}$, L339$^{8.57}$), (N69$^{2.40}$, D331$^{8.49}$), (I72$^{2.43}$, P323$^{7.50}$), (V87$^{2.58}$, Y316$^{7.43}$), (G90$^{2.61}$, Y316$^{7.43}$) |

**Table S4.** Residue pairs with significant change in average nonbonding interaction energies ($|\Delta\langle E_{XY}\rangle| > 30$ kcal/mol) within the TM region

| Residue X | Residue Y | $\Delta\langle E_{XY}\rangle$(kcal/mol) | $\langle E_{XY}\rangle^{\mathrm{B2ARP}}$(kcal/mol) | $\langle E_{XY}\rangle^{\mathrm{B2AR}}$(kcal/mol) |
|---|---|---|---|---|
| T360 | K263 | -190.5259 | -190.4959 | 0.03 |
| S411 | K273 | -186.908 | -186.8424 | 0.0655 |
| S411 | K270 | -185.7244 | -185.7196 | 47 |
| S407 | R259 | -169.5496 | -169.5852 | -0.0356 |
| S407 | K263 | -137.7756 | -137.8207 | -0.045 |
| T360 | R239 | -123.6843 | -123.7147 | -0.0304 |
| S364 | R239 | -79.5447 | -80.9604 | -1.4158 |
| R344 | E338 | -76.9976 | -103.2363 | -26.2387 |
| S411 | R333 | -73.3843 | -73.9306 | -0.5462 |
| T360 | R259 | -71.6907 | -71.6884 | 24 |
| D386 | R63 | -71.6902 | -90.1013 | -18.4112 |
| D410 | K273 | -71.3676 | -94.3716 | -23.004 |
| S411 | R131 | -62.5043 | -62.5113 | -7 |
| S411 | K267 | -62.2386 | -62.2468 | -82 |
| S407 | R239 | -60.9493 | -61.7604 | -0.8111 |
| S411 | R328 | -53.6624 | -54.0183 | -0.3559 |
| S401 | R239 | -53.549 | -53.723 | -0.174 |
| S407 | R260 | -52.2623 | -52.2671 | -48 |
| S411 | K263 | -48.6043 | -48.5287 | 0.0756 |
| T360 | K273 | -48.1305 | -48.0785 | 0.052 |
| S407 | K273 | -44.8721 | -44.8612 | 0.011 |
| S411 | R239 | -44.5524 | -44.3959 | 0.1565 |
| T360 | K267 | -44.3611 | -44.316 | 0.0451 |
| S396 | R239 | -43.5307 | -43.6234 | -0.0927 |
| T360 | K270 | -41.513 | -41.4361 | 0.0769 |
| S364 | K263 | -40.1454 | -40.2976 | -0.1523 |
| S401 | K263 | -39.4523 | -39.5853 | -0.133 |
| T360 | R260 | -38.7639 | -38.7513 | 0.0126 |
| S411 | R260 | -37.4722 | -37.4333 | 0.0389 |
| S411 | R259 | -37.1007 | -37.0416 | 0.0591 |
| S411 | K227 | -36.963 | -36.953 | 0.0101 |
| S401 | R333 | -35.4145 | -35.6522 | -0.2378 |
| S396 | R333 | -35.1101 | -35.3009 | -0.1908 |
| S407 | K270 | -35.0291 | -35.4496 | -0.4205 |
| T360 | R333 | -34.9838 | -34.9256 | 0.0582 |
| S407 | K267 | -34.9145 | -35.7126 | -0.7981 |
| S411 | K232 | -34.6885 | -34.5077 | 0.1809 |
| S364 | K267 | -34.1036 | -34.2933 | -0.1896 |
| S364 | R333 | -33.7934 | -33.7643 | 0.0292 |
| S364 | K270 | -33.2129 | -33.2692 | -0.0564 |
| T360 | K235 | -32.8468 | -32.8654 | -0.0186 |
| S364 | K235 | -32.7945 | -32.9907 | -0.1961 |

| | | | | |
|---|---|---|---|---|
| S401 | K273 | -32.4551 | -32.6248 | -0.1697 |
| S364 | K273 | -32.3713 | -32.4327 | -0.0615 |
| S407 | K227 | -32.1366 | -32.2234 | -0.0868 |
| S396 | K263 | -31.8277 | -31.9031 | -0.0753 |
| S396 | R63 | -31.7522 | -31.8999 | -0.1478 |
| S407 | R333 | -31.6779 | -31.4073 | 0.2706 |
| S401 | K270 | -31.5909 | -31.8609 | -0.2699 |
| T360 | K227 | -30.7224 | -30.7161 | 63 |
| S364 | R63 | -30.5342 | -30.4946 | 0.0396 |
| E373 | K235 | -30.4124 | -42.0853 | -11.6729 |
| S401 | K267 | -30.3633 | -30.5441 | -0.1807 |
| S411 | K235 | -30.3369 | -30.1735 | 0.1634 |
| S396 | K270 | -30.2536 | -30.3739 | -0.1203 |
| S364 | D331 | 31.0753 | 31.0502 | -0.0251 |
| S396 | D331 | 31.5675 | 31.8501 | 0.2826 |
| S411 | E338 | 32.0389 | 32.0547 | 0.0158 |
| S401 | E237 | 32.1235 | 32.3186 | 0.1951 |
| S411 | E225 | 34.3279 | 34.1691 | -0.1588 |
| T360 | E268 | 35.9734 | 35.964 | -93 |
| S407 | E237 | 36.1318 | 36.4314 | 0.2996 |
| S364 | E338 | 36.5345 | 36.4982 | -0.0363 |
| S407 | E268 | 36.8234 | 36.9418 | 0.1184 |
| S364 | E237 | 37.3173 | 37.0872 | -0.2301 |
| S401 | E338 | 38.0238 | 38.0958 | 0.0721 |
| R404 | K263 | 40.3743 | 54.5105 | 14.1362 |
| R343 | E338 | 41.8836 | -28.8404 | -70.724 |
| S411 | E268 | 43.7863 | 43.4904 | -0.2959 |
| T360 | E237 | 47.0408 | 46.9584 | -0.0823 |
| S396 | E338 | 48.4135 | 48.5037 | 0.0902 |
| S411 | E237 | 56.4514 | 56.3597 | -0.0917 |
| S411 | D331 | 58.6661 | 54.1505 | -4.5156 |

**Table S5.** Pairwise change in net transfer entropies. Residue pairs with the highest $\Delta_{\mathbf{Net}}\mathbf{TE}_{XY}$ are shown.

| **Donor (X)** | **Receiver (Y)** | $\Delta_{\mathbf{Net}}\mathbf{TE}_{XY}$ | $\mathbf{NetTE}_{XY}^{B2ARP}$ | $\mathbf{NetTE}_{XY}^{B2AR}$ |
|---|---|---|---|---|
| A349 | $L45^{1.44}$ | 0.0182 | 0.0169 | -0.0013 |
| G351 | $A46^{1.45}$ | 0.0182 | 0.007 | -0.0112 |
| L381 | $A78^{2.49}$ | 0.0126 | 0.0082 | -0.0044 |
| F387 | $A78^{2.49}$ | -0.026 | 0.0063 | 0.0323 |
| V388 | $A78^{2.49}$ | -0.0289 | 0.0013 | 0.0303 |
| L381 | $D79^{2.50}$ | 0.0096 | 0.005 | -0.0045 |
| V388 | $D79^{2.50}$ | -0.027 | -0.0047 | 0.0223 |
| H390 | $D79^{2.50}$ | -0.0276 | -0.0059 | 0.0217 |
| T384 | $V81^{2.52}$ | -0.0279 | -0.0029 | 0.0251 |
| Q370 | $M82^{2.53}$ | 0.0202 | 0.0124 | -0.0078 |
| V388 | $M82^{2.53}$ | -0.0273 | 0.001 | 0.0284 |
| I399 | $M82^{2.53}$ | 0.0203 | 0.0103 | -0.01 |
| V388 | $A85^{2.56}$ | -0.026 | -0.0056 | 0.0204 |
| T393 | $V117^{3.36}$ | -0.0315 | 0.0001 | 0.0316 |
| E369 | $T118^{3.37}$ | 0.0177 | 0.0143 | -0.0034 |
| D386 | $T118^{3.37}$ | -0.0312 | 0.0008 | 0.0319 |
| F387 | $T118^{3.37}$ | -0.0379 | 0.0037 | 0.0416 |
| V388 | $T118^{3.37}$ | -0.0379 | -0.0036 | 0.0343 |
| T393 | $T118^{3.37}$ | -0.0265 | 0.0106 | 0.0371 |
| D386 | $A119^{3.38}$ | -0.029 | 0.0011 | 0.0301 |
| F387 | $A119^{3.38}$ | -0.0258 | 0.0075 | 0.0333 |
| V388 | $A119^{3.38}$ | -0.0362 | -0.0021 | 0.0342 |
| H390 | $A119^{3.38}$ | -0.0289 | 0.0057 | 0.0345 |
| H390 | $S120^{3.39}$ | -0.0313 | -0.0028 | 0.0284 |
| Q391 | $S120^{3.39}$ | -0.0262 | -0.0019 | 0.0244 |
| T408 | $S120^{3.39}$ | 0.0194 | 0.0047 | -0.0147 |
| L342 | $T123^{3.42}$ | 0.0205 | 0.0046 | -0.0159 |
| E369 | $T123^{3.42}$ | 0.019 | 0.0144 | -0.0046 |
| D386 | $T123^{3.42}$ | -0.0329 | 0.0014 | 0.0344 |
| V388 | $T123^{3.42}$ | -0.0331 | -0.0029 | 0.0302 |
| Q370 | $M156^{4.48}$ | 0.0134 | 0.0086 | -0.0048 |
| S355 | $M215^{5.54}$ | 0.0197 | 0.0126 | -0.0071 |
| T360 | $M215^{5.54}$ | 0.0111 | 0.0125 | 0.0013 |
| S364 | $M215^{5.54}$ | 0.0044 | 0.0064 | 0.002 |
| E369 | $M215^{5.54}$ | 0.0067 | 0.0036 | -0.0031 |
| Q370 | $M215^{5.54}$ | 0.0034 | 0.0038 | 0.0003 |
| S396 | $M215^{5.54}$ | 0.0017 | 0.0017 | 0 |
| S401 | $M215^{5.54}$ | 0.0076 | 0.0076 | 0 |
| E371 | $F217^{5.56}$ | 0.0039 | 0.0029 | -0.001 |
| E369 | $V218^{5.57}$ | 0.0002 | 0.0021 | 0.0019 |
| E371 | $V218^{5.57}$ | 0.0127 | 0.0079 | -0.0048 |

| | | | | |
|---|---|---|---|---|
| E373 | V218$^{5.57}$ | 0.0061 | 0.0067 | 0.0006 |
| N398 | V218$^{5.57}$ | 0.008 | 0.011 | 0.003 |
| S401 | Y219$^{5.58}$ | -0.013 | -0.0032 | 0.0098 |
| E369 | S220$^{5.59}$ | 0.0042 | 0.009 | 0.0048 |
| Q370 | S220$^{5.59}$ | 0.0046 | 0.0072 | 0.0026 |
| E371 | S220$^{5.59}$ | 0.0127 | 0.0136 | 0.0008 |
| T360 | M279$^{6.41}$ | 0.0081 | 0.0081 | 0 |
| S364 | M279$^{6.41}$ | 0.0087 | 0.0087 | 0 |
| S396 | M279$^{6.41}$ | 0.0015 | 0.0015 | 0 |
| S401 | M279$^{6.41}$ | 0.0011 | 0.0011 | 0 |
| S411 | M279$^{6.41}$ | 0.0011 | 0.0058 | 0.0046 |
| T360 | F282$^{6.44}$ | 0.008 | 0.008 | 0 |
| S364 | F282$^{6.44}$ | 0.0062 | 0.0062 | 0 |
| S396 | F282$^{6.44}$ | 0.0036 | 0.0036 | 0 |
| S411 | F282$^{6.44}$ | 0.0033 | 0.0033 | 0 |
| E369 | C285$^{6.47}$ | 0.0167 | 0.0093 | -0.0074 |
| E369 | E62 | 0.0081 | 0.0077 | -0.0004 |
| Q370 | R63 | 0.0141 | 0.0094 | -0.0046 |
| E371 | R63 | 0.0095 | 0.0079 | -0.0016 |
| E369 | V67$^{2.38}$ | 0.0251 | 0.0227 | -0.0024 |
| Q370 | T68$^{2.39}$ | 0.0175 | 0.0132 | -0.0043 |
| Q370 | R131$^{3.50}$ | 0.019 | 0.0094 | -0.0096 |
| E369 | Y132$^{3.51}$ | 0.0043 | 0.0042 | -0.0001 |
| Q370 | Y132$^{3.51}$ | 0.0041 | 0.003 | -0.0011 |
| E371 | Y132$^{3.51}$ | 0.0111 | 0.0103 | -0.0008 |
| R343 | P138 | -0.018 | -0.0053 | 0.0127 |
| Y354 | K147$^{4.39}$ | 0.0225 | 0.0225 | 0 |
| L342 | N148$^{4.40}$ | 0.0243 | 0.0132 | -0.011 |
| D410 | K149$^{4.41}$ | -0.0199 | -0.0098 | 0.0101 |
| N398 | A150$^{4.42}$ | 0.0065 | -0.0021 | -0.0086 |
| K348 | R151$^{4.43}$ | 0.0224 | 0.0145 | -0.0079 |
| Q370 | R151$^{4.43}$ | 0.0243 | 0.0125 | -0.0118 |
| N398 | R151$^{4.43}$ | 0.0067 | 0.0003 | -0.0065 |
| E369 | V152$^{4.44}$ | 0.0184 | 0.014 | -0.0043 |
| Q370 | V152$^{4.44}$ | 0.0184 | 0.0137 | -0.0046 |
| E369 | I153$^{4.45}$ | 0.0015 | 0.0003 | -0.0011 |
| S356 | I154$^{4.46}$ | 0.0217 | 0.0217 | 0 |
| Q363 | I154$^{4.46}$ | 0.0197 | 0.0197 | 0 |
| S364 | I154$^{4.46}$ | 0.0184 | 0.0184 | 0 |
| H390 | L230$^{5.69}$ | 0.0136 | 0.0136 | 0 |
| T408 | L230$^{5.69}$ | 0.0113 | 0.0075 | -0.0038 |
| S411 | E237 | -0.0163 | -0.0127 | 0.0037 |
| S346 | R239 | -0.0168 | -0.0106 | 0.0062 |
| G353 | R239 | 0.0017 | 0.0049 | 0.0032 |
| R344 | Q247 | -0.0171 | -0.0137 | 0.0035 |
| K348 | G252 | 0.0125 | 0.0047 | -0.0078 |

| | | | | |
|---|---|---|---|---|
| S411 | H256 | -0.0184 | -0.0124 | 0.006 |
| N409 | $C265^{6.27}$ | -0.0203 | -0.0085 | 0.0118 |
| E369 | $K267^{6.29}$ | 0.0166 | 0.0072 | -0.0094 |
| G392 | $E268^{6.30}$ | 0.0127 | 0.0127 | 0 |
| G383 | $N322^{7.49}$ | -0.0232 | -0.0082 | 0.015 |
| T384 | $N322^{7.49}$ | -0.0268 | -0.0077 | 0.0191 |
| P395 | $N322^{7.49}$ | -0.017 | -0.0017 | 0.0153 |
| S396 | $N322^{7.49}$ | -0.0149 | -0.0096 | 0.0053 |
| N352 | $P323^{7.50}$ | 0.0164 | 0.0131 | -0.0033 |
| T384 | $P323^{7.50}$ | -0.0164 | 0.0072 | 0.0236 |
| E385 | $P323^{7.50}$ | -0.022 | -0.0028 | 0.0192 |
| D386 | $P323^{7.50}$ | -0.0196 | 0.0025 | 0.0221 |
| N409 | $P323^{7.50}$ | -0.0294 | -0.0089 | 0.0205 |
| D410 | $P323^{7.50}$ | -0.0164 | 0.0002 | 0.0166 |
| D410 | $L324^{7.51}$ | -0.0198 | -0.0063 | 0.0134 |
| R344 | $I325^{7.52}$ | -0.0212 | -0.0127 | 0.0085 |
| D410 | $I325^{7.52}$ | -0.016 | -0.0147 | 0.0012 |
| L377 | $Y326^{7.53}$ | 0.0172 | 0.0172 | 0 |
| G353 | $C327^{7.54}$ | 0.014 | 0.0044 | -0.0096 |
| G353 | $F332^{8.50}$ | 0.0149 | 0.01 | -0.005 |
| G392 | $F332^{8.50}$ | 0.015 | 0.015 | 0 |
| A349 | $R333^{8.51}$ | 0.0142 | 0.0121 | -0.0021 |
| D386 | $R333^{8.51}$ | -0.0182 | -0.0042 | 0.014 |
| H390 | $I334^{8.52}$ | 0.017 | 0.017 | 0 |
| S346 | $F336^{8.54}$ | 0.0168 | 0.0091 | -0.0077 |
| G392 | $Q337^{8.55}$ | 0.016 | 0.016 | 0 |
| E373 | $L339^{8.57}$ | 0.0144 | 0.0144 | 0 |
| $N322^{7.49}$ | $V48^{1.47}$ | 0.0245 | 0.0087 | -0.0158 |
| $C327^{7.54}$ | $D79^{2.50}$ | -0.0225 | -0.0031 | 0.0194 |
| $P323^{7.50}$ | $L80^{2.51}$ | 0.0262 | 0.0055 | -0.0207 |
| $L324^{7.51}$ | $L80^{2.51}$ | 0.0228 | 0.0081 | -0.0147 |
| $V152^{4.44}$ | $V81^{2.52}$ | -0.02 | -0.0082 | 0.0119 |
| $C327^{7.54}$ | $V81^{2.52}$ | -0.0263 | -0.0051 | 0.0213 |
| F240 | $T118^{3.37}$ | 0.0237 | 0.0128 | -0.0109 |
| $T66^{2.37}$ | $I121^{3.40}$ | 0.0231 | 0.001 | -0.022 |
| $N69^{2.40}$ | $L124^{3.43}$ | 0.0262 | 0.0037 | -0.0225 |
| $I154^{4.46}$ | $V157^{4.49}$ | -0.0188 | -0.0132 | 0.0056 |
| $V152^{4.44}$ | $V213^{5.52}$ | -0.0235 | -0.0102 | 0.0134 |
| $F336^{8.54}$ | $V213^{5.52}$ | -0.0284 | -0.0206 | 0.0078 |
| $N148^{4.40}$ | $I214^{5.53}$ | 0.0137 | 0.0089 | -0.0048 |
| $L230^{5.69}$ | $I214^{5.53}$ | 0.0083 | 0.0093 | 0.0011 |
| $K235^{5.74}$ | $I214^{5.53}$ | 0.0108 | 0.0125 | 0.0018 |
| $F264^{6.26}$ | $I214^{5.53}$ | -0.0126 | -0.0029 | 0.0097 |
| $C265^{6.27}$ | $I214^{5.53}$ | 0.0135 | 0.0072 | -0.0062 |
| $R131^{3.50}$ | $M215^{5.54}$ | 0.0102 | 0.0044 | -0.0057 |
| $R228^{5.67}$ | $M215^{5.54}$ | 0.0083 | 0.0009 | -0.0075 |

| | | | | |
|---|---|---|---|---|
| K232$^{5.71}$ | M215$^{5.54}$ | 0.0118 | 0.0105 | -0.0013 |
| D234$^{5.73}$ | M215$^{5.54}$ | 0.0131 | 0.014 | 0.001 |
| K235$^{5.74}$ | M215$^{5.54}$ | 0.0081 | 0.0094 | 0.0013 |
| S262$^{6.24}$ | M215$^{5.54}$ | 0.0133 | 0.0101 | -0.0032 |
| K263$^{6.25}$ | M215$^{5.54}$ | 0.0115 | 0.0102 | -0.0013 |
| C265$^{6.27}$ | M215$^{5.54}$ | 0.0149 | 0.0107 | -0.0042 |
| K232$^{5.71}$ | V216$^{5.55}$ | 0.0104 | 0.0094 | -0.001 |
| I233$^{5.72}$ | V216$^{5.55}$ | 0.0163 | 0.0138 | -0.0025 |
| I135$^{3.54}$ | F217$^{5.56}$ | -0.0198 | -0.0189 | 0.0009 |
| V152$^{4.44}$ | F217$^{5.56}$ | -0.0266 | -0.0189 | 0.0077 |
| E268$^{6.30}$ | F217$^{5.56}$ | -0.0142 | -0.0094 | 0.0048 |
| R228$^{5.67}$ | V218$^{5.57}$ | 0.0108 | 0.0089 | -0.0019 |
| K232$^{5.71}$ | V218$^{5.57}$ | 0.0111 | 0.0086 | -0.0025 |
| F240 | V218$^{5.57}$ | 0.0244 | 0.0092 | -0.0153 |
| C265$^{6.27}$ | V218$^{5.57}$ | 0.0116 | 0.0072 | -0.0044 |
| R131$^{3.50}$ | Y219$^{5.58}$ | -0.0214 | -0.0206 | 0.0008 |
| L145 | Y219$^{5.58}$ | -0.0237 | -0.0078 | 0.0158 |
| R151$^{4.43}$ | Y219$^{5.58}$ | -0.0187 | -0.0129 | 0.0058 |
| V152$^{4.44}$ | Y219$^{5.58}$ | -0.0206 | -0.0148 | 0.0058 |
| I154$^{4.46}$ | Y219$^{5.58}$ | -0.0184 | -0.0111 | 0.0073 |
| K235$^{5.74}$ | Y219$^{5.58}$ | 0.0151 | 0.0114 | -0.0038 |
| E62 | Y219$^{5.58}$ | -0.0249 | -0.0185 | 0.0064 |
| L64 | Y219$^{5.58}$ | -0.0227 | -0.0152 | 0.0076 |
| V67$^{2.38}$ | Y219$^{5.58}$ | -0.0174 | -0.0105 | 0.0069 |
| K232$^{5.71}$ | M279$^{6.41}$ | 0.0123 | 0.0097 | -0.0026 |
| C265$^{6.27}$ | M279$^{6.41}$ | 0.0138 | 0.0064 | -0.0074 |
| K232$^{5.71}$ | G280$^{6.42}$ | 0.0091 | 0.0098 | 0.0007 |
| I233$^{5.72}$ | G280$^{6.42}$ | 0.0126 | 0.0165 | 0.0039 |
| D234$^{5.73}$ | G280$^{6.42}$ | 0.0126 | 0.0067 | -0.0059 |
| K263$^{6.25}$ | G280$^{6.42}$ | 0.0083 | 0.0103 | 0.002 |
| F264$^{6.26}$ | T281$^{6.43}$ | -0.012 | -0.0018 | 0.0102 |
| K267$^{6.29}$ | T281$^{6.43}$ | 0.0107 | 0.0028 | -0.008 |
| R228$^{5.67}$ | F282$^{6.44}$ | 0.0108 | 0.004 | -0.0068 |
| Q231$^{5.70}$ | F282$^{6.44}$ | 0.0088 | 0.0078 | -0.001 |
| K232$^{5.71}$ | F282$^{6.44}$ | 0.0122 | 0.0074 | -0.0048 |
| I233$^{5.72}$ | F282$^{6.44}$ | 0.0134 | 0.0074 | -0.006 |
| D234$^{5.73}$ | F282$^{6.44}$ | 0.0186 | 0.0102 | -0.0084 |
| K235$^{5.74}$ | F282$^{6.44}$ | 0.0099 | 0.0081 | -0.0018 |
| C265$^{6.27}$ | F282$^{6.44}$ | 0.011 | 0.0017 | -0.0093 |
| R228$^{5.67}$ | T283$^{6.45}$ | 0.0086 | 0.0046 | -0.004 |
| L230$^{5.69}$ | T283$^{6.45}$ | -0.0107 | -0.0002 | 0.0105 |
| Q231$^{5.70}$ | T283$^{6.45}$ | 0.0082 | 0.0071 | -0.0011 |
| K235$^{5.74}$ | T283$^{6.45}$ | 0.0114 | 0.004 | -0.0074 |
| C265$^{6.27}$ | T283$^{6.45}$ | 0.0084 | 0.0063 | -0.0022 |
| L266$^{6.28}$ | T283$^{6.45}$ | 0.0121 | 0.0049 | -0.0072 |
| R151$^{4.43}$ | L284$^{6.46}$ | -0.0177 | -0.0131 | 0.0045 |

| | | | | |
|---|---|---|---|---|
| E268$^{6.30}$ | L284$^{6.46}$ | -0.0109 | -0.0031 | 0.0078 |
| H269$^{6.31}$ | L284$^{6.46}$ | -0.0119 | -0.0131 | -0.0012 |
| R151$^{4.43}$ | S319$^{7.46}$ | -0.0221 | -0.006 | 0.0161 |
| V152$^{4.44}$ | S319$^{7.46}$ | -0.0228 | -0.0124 | 0.0104 |
| I154$^{4.46}$ | S319$^{7.46}$ | -0.0226 | -0.0084 | 0.0142 |
| L64 | S319$^{7.46}$ | -0.0208 | -0.0086 | 0.0122 |
| A349 | L45$^{1.44}$ | 0.0182 | 0.0169 | -0.0013 |
| G351 | A46$^{1.45}$ | 0.0182 | 0.007 | -0.0112 |
| L381 | A78$^{2.49}$ | 0.0126 | 0.0082 | -0.0044 |
| F387 | A78$^{2.49}$ | -0.026 | 0.0063 | 0.0323 |
| V388 | A78$^{2.49}$ | -0.0289 | 0.0013 | 0.0303 |
| L381 | D79$^{2.50}$ | 0.0096 | 0.005 | -0.0045 |
| V388 | D79$^{2.50}$ | -0.027 | -0.0047 | 0.0223 |
| H390 | D79$^{2.50}$ | -0.0276 | -0.0059 | 0.0217 |
| T384 | V81$^{2.52}$ | -0.0279 | -0.0029 | 0.0251 |
| Q370 | M82$^{2.53}$ | 0.0202 | 0.0124 | -0.0078 |
| V388 | M82$^{2.53}$ | -0.0273 | 0.001 | 0.0284 |
| I399 | M82$^{2.53}$ | 0.0203 | 0.0103 | -0.01 |
| V388 | A85$^{2.56}$ | -0.026 | -0.0056 | 0.0204 |
| T393 | V117$^{3.36}$ | -0.0315 | 0.0001 | 0.0316 |
| E369 | T118$^{3.37}$ | 0.0177 | 0.0143 | -0.0034 |
| D386 | T118$^{3.37}$ | -0.0312 | 0.0008 | 0.0319 |
| F387 | T118$^{3.37}$ | -0.0379 | 0.0037 | 0.0416 |
| V388 | T118$^{3.37}$ | -0.0379 | -0.0036 | 0.0343 |
| T393 | T118$^{3.37}$ | -0.0265 | 0.0106 | 0.0371 |
| D386 | A119$^{3.38}$ | -0.029 | 0.0011 | 0.0301 |
| F387 | A119$^{3.38}$ | -0.0258 | 0.0075 | 0.0333 |
| V388 | A119$^{3.38}$ | -0.0362 | -0.0021 | 0.0342 |
| H390 | A119$^{3.38}$ | -0.0289 | 0.0057 | 0.0345 |
| H390 | S120$^{3.39}$ | -0.0313 | -0.0028 | 0.0284 |
| Q391 | S120$^{3.39}$ | -0.0262 | -0.0019 | 0.0244 |
| T408 | S120$^{3.39}$ | 0.0194 | 0.0047 | -0.0147 |
| L342 | T123$^{3.42}$ | 0.0205 | 0.0046 | -0.0159 |
| E369 | T123$^{3.42}$ | 0.019 | 0.0144 | -0.0046 |
| D386 | T123$^{3.42}$ | -0.0329 | 0.0014 | 0.0344 |
| V388 | T123$^{3.42}$ | -0.0331 | -0.0029 | 0.0302 |
| Q370 | M156$^{4.48}$ | 0.0134 | 0.0086 | -0.0048 |
| S355 | M215$^{5.54}$ | 0.0197 | 0.0126 | -0.0071 |
| T360 | M215$^{5.54}$ | 0.0111 | 0.0125 | 0.0013 |
| S364 | M215$^{5.54}$ | 0.0044 | 0.0064 | 0.002 |
| E369 | M215$^{5.54}$ | 0.0067 | 0.0036 | -0.0031 |
| Q370 | M215$^{5.54}$ | 0.0034 | 0.0038 | 0.0003 |
| S396 | M215$^{5.54}$ | 0.0017 | 0.0017 | 0 |
| S401 | M215$^{5.54}$ | 0.0076 | 0.0076 | 0 |
| E371 | F217$^{5.56}$ | 0.0039 | 0.0029 | -0.001 |
| E369 | V218$^{5.57}$ | 0.0002 | 0.0021 | 0.0019 |

| | | | | |
|---|---|---|---|---|
| E371 | V218$^{5.57}$ | 0.0127 | 0.0079 | -0.0048 |
| E373 | V218$^{5.57}$ | 0.0061 | 0.0067 | 0.0006 |
| N398 | V218$^{5.57}$ | 0.008 | 0.011 | 0.003 |
| S401 | Y219$^{5.58}$ | -0.013 | -0.0032 | 0.0098 |
| E369 | S220$^{5.59}$ | 0.0042 | 0.009 | 0.0048 |
| Q370 | S220$^{5.59}$ | 0.0046 | 0.0072 | 0.0026 |

**Table S6.** Residue pairs with significant change in average nonbonding interaction energies ($|\Delta\langle E_{XY}\rangle| > 8$ kcal/mol) within the TM region

| Residue X | Residue Y | $\Delta\langle E_{XY}\rangle$(kcal/mol) | $\langle E_{XY}\rangle^{B2ARP}$(kcal/mol) | $\langle E_{XY}\rangle^{B2AR}$(kcal/mol) |
|---|---|---|---|---|
| E237 | $K267^{6.29}$ | -64.7102 | -100.1314 | -35.4212 |
| R239 | $K267^{6.29}$ | -30.4955 | 23.8718 | 54.3673 |
| $E225^{5.64}$ | $K232^{5.71}$ | -28.5082 | -71.7547 | -43.2465 |
| $R228^{5.67}$ | E249 | -25.0886 | -38.6035 | -13.515 |
| $K232^{5.71}$ | $D234^{5.73}$ | -20.2885 | -46.0654 | -25.7769 |
| $K227^{5.66}$ | E249 | -19.0672 | -34.9082 | -15.8411 |
| R260 | $E268^{6.30}$ | -17.3023 | -33.8304 | -16.5281 |
| R253 | R259 | -17.1111 | 12.5338 | 29.6449 |
| G257 | R260 | -16.5101 | -15.2398 | 1.2703 |
| $K227^{5.66}$ | $E268^{6.30}$ | -16.2026 | -96.6421 | -80.4395 |
| $K232^{5.71}$ | E237 | -15.6252 | -37.8356 | -22.2104 |
| $K232^{5.71}$ | $K235^{5.74}$ | -13.1413 | 23.0124 | 36.1536 |
| $E225^{5.64}$ | $K267^{6.29}$ | -12.7309 | -33.2373 | -20.5064 |
| $D234^{5.73}$ | $E268^{6.30}$ | -12.3464 | 18.083 | 30.4293 |
| $R228^{5.67}$ | $K235^{5.74}$ | -11.8844 | 15.7513 | 27.6357 |
| L258 | R260 | -11.0693 | -10.0229 | 1.0464 |
| $D234^{5.73}$ | $K235^{5.74}$ | -10.8716 | -99.6353 | -88.7637 |
| E237 | $K270^{6.32}$ | -10.7567 | -32.2196 | -21.4629 |
| E237 | $K273^{6.35}$ | -10.1047 | -24.9786 | -14.8739 |
| $D234^{5.73}$ | E237 | -9.3867 | 27.2902 | 36.677 |
| $K235^{5.74}$ | E237 | -9.2494 | -31.8794 | -22.63 |
| $K227^{5.66}$ | R259 | -8.8074 | 16.898 | 25.7053 |
| R260 | $H269^{6.31}$ | -8.6619 | -8.8334 | -0.1716 |
| $K232^{5.71}$ | E249 | -8.6508 | -19.6116 | -10.9608 |
| R260 | S261 | -8.0581 | -66.3639 | -58.3058 |
| $Q224^{5.63}$ | $K227^{5.66}$ | 8.0338 | -9.5446 | -17.5784 |
| D251 | R253 | 8.5102 | -34.5248 | -43.035 |
| $K235^{5.74}$ | $E268^{6.30}$ | 9.1438 | -15.4448 | -24.5886 |
| $E225^{5.64}$ | E249 | 9.3124 | 20.934 | 11.6216 |
| $E225^{5.64}$ | E237 | 9.8273 | 26.2567 | 16.4294 |
| H256 | R260 | 10.0167 | 4.0023 | -6.0144 |
| E249 | $E268^{6.30}$ | 11.5609 | 28.3795 | 16.8185 |
| R239 | R259 | 12.1066 | 24.6364 | 12.5298 |
| $K227^{5.66}$ | R253 | 12.6492 | 27.9948 | 15.3456 |
| $D234^{5.73}$ | $K270^{6.32}$ | 13.6866 | -16.0501 | -29.7367 |
| $K270^{6.32}$ | $K273^{6.35}$ | 13.9141 | 56.7131 | 42.799 |
| $K232^{5.71}$ | $K267^{6.29}$ | 14.3198 | 37.7641 | 23.4443 |
| E249 | R259 | 15.4934 | -14.6436 | -30.137 |
| $K227^{5.66}$ | R260 | 16.6144 | 31.8442 | 15.2298 |
| $E225^{5.64}$ | $R228^{5.67}$ | 17.0634 | -73.4029 | -90.4663 |
| R239 | $K263^{6.25}$ | 28.4529 | 49.3567 | 20.9039 |
| R259 | $K263^{6.25}$ | 29.3119 | 47.2911 | 17.9792 |

| | | | | |
|---|---|---|---|---|
| E237 | R239 | 36.1869 | -28.1254 | -64.3122 |
| D251 | R259 | 64.6578 | -14.7618 | -79.4196 |
| $D234^{5.73}$ | R239 | 67.1047 | -16.0467 | -83.1515 |
| $D234^{5.73}$ | $K267^{6.29}$ | 81.5366 | -26.453 | -107.9896 |
| $D234^{5.73}$ | K140 | -15.161 | -28.1637 | -13.0027 |
| $K267^{6.29}$ | K140 | 8.4695 | 21.0487 | 12.5792 |
| $K273^{6.35}$ | $R131^{3.50}$ | 10.9765 | 28.4014 | 17.4249 |
| $K270^{6.32}$ | $R131^{3.50}$ | 22.9212 | 42.5172 | 19.596 |
| $D331^{8.49}$ | $K270^{6.32}$ | -14.0891 | -30.8478 | -16.7587 |
| $K305^{7.32}$ | D192 | -12.1466 | -98.6639 | -86.5173 |
| $E306^{7.33}$ | K97 | -9.2844 | -42.5168 | -33.2324 |
| $D331^{8.49}$ | $K273^{6.35}$ | -8.5135 | -22.5921 | -14.0786 |
| $D331^{8.49}$ | $R131^{3.50}$ | -8.1444 | -40.3479 | -32.2034 |
| $R333^{8.51}$ | $K270^{6.32}$ | 8.8694 | 31.6713 | 22.8018 |
| $Y316^{7.43}$ | $D113^{3.32}$ | 10.0278 | -5.3002 | -15.328 |
| $R333^{8.51}$ | $K273^{6.35}$ | 13.8473 | 33.5044 | 19.657 |
| $D331^{8.49}$ | R63 | 21.8999 | -23.9172 | -45.8171 |
| $E338^{8.56}$ | R63 | 23.0424 | -22.4659 | -45.5083 |

**Table S7.** Residue pairs with significant change in average van der Waal interaction energies ($|\Delta\langle E_{XY}^{\mathbf{vdw}}\rangle| > 1$ kcal/mol) within the TM region

| **Residue X** | **Residue Y** | $\Delta\langle E_{XY}^{\mathbf{vdw}}\rangle$**(kcal/mol)** | $\langle E_{XY}^{\mathbf{vdw}}\rangle^{\mathbf{B2ARP}}$**(kcal/mol)** | $\langle E_{XY}^{\mathbf{vdw}}\rangle^{\mathbf{B2AR}}$**(kcal/mol)** |
|---|---|---|---|---|
| A349 | $L45^{1.44}$ | 0.0182 | 0.0169 | -0.0013 |
| G351 | $A46^{1.45}$ | 0.0182 | 0.007 | -0.0112 |
| L381 | $A78^{2.49}$ | 0.0126 | 0.0082 | -0.0044 |
| F387 | $A78^{2.49}$ | -0.026 | 0.0063 | 0.0323 |
| V388 | $A78^{2.49}$ | -0.0289 | 0.0013 | 0.0303 |
| L381 | $D79^{2.50}$ | 0.0096 | 0.005 | -0.0045 |
| V388 | $D79^{2.50}$ | -0.027 | -0.0047 | 0.0223 |
| H390 | $D79^{2.50}$ | -0.0276 | -0.0059 | 0.0217 |
| T384 | $V81^{2.52}$ | -0.0279 | -0.0029 | 0.0251 |
| Q370 | $M82^{2.53}$ | 0.0202 | 0.0124 | -0.0078 |
| V388 | $M82^{2.53}$ | -0.0273 | 0.001 | 0.0284 |
| I399 | $M82^{2.53}$ | 0.0203 | 0.0103 | -0.01 |
| V388 | $A85^{2.56}$ | -0.026 | -0.0056 | 0.0204 |
| T393 | $V117^{3.36}$ | -0.0315 | 0.0001 | 0.0316 |
| E369 | $T118^{3.37}$ | 0.0177 | 0.0143 | -0.0034 |
| D386 | $T118^{3.37}$ | -0.0312 | 0.0008 | 0.0319 |
| F387 | $T118^{3.37}$ | -0.0379 | 0.0037 | 0.0416 |
| V388 | $T118^{3.37}$ | -0.0379 | -0.0036 | 0.0343 |
| T393 | $T118^{3.37}$ | -0.0265 | 0.0106 | 0.0371 |
| D386 | $A119^{3.38}$ | -0.029 | 0.0011 | 0.0301 |
| F387 | $A119^{3.38}$ | -0.0258 | 0.0075 | 0.0333 |
| V388 | $A119^{3.38}$ | -0.0362 | -0.0021 | 0.0342 |
| H390 | $A119^{3.38}$ | -0.0289 | 0.0057 | 0.0345 |
| H390 | $S120^{3.39}$ | -0.0313 | -0.0028 | 0.0284 |
| Q391 | $S120^{3.39}$ | -0.0262 | -0.0019 | 0.0244 |
| T408 | $S120^{3.39}$ | 0.0194 | 0.0047 | -0.0147 |
| L342 | $T123^{3.42}$ | 0.0205 | 0.0046 | -0.0159 |
| E369 | $T123^{3.42}$ | 0.019 | 0.0144 | -0.0046 |
| D386 | $T123^{3.42}$ | -0.0329 | 0.0014 | 0.0344 |
| V388 | $T123^{3.42}$ | -0.0331 | -0.0029 | 0.0302 |
| Q370 | $M156^{4.48}$ | 0.0134 | 0.0086 | -0.0048 |
| S355 | $M215^{5.54}$ | 0.0197 | 0.0126 | -0.0071 |
| T360 | $M215^{5.54}$ | 0.0111 | 0.0125 | 0.0013 |
| S364 | $M215^{5.54}$ | 0.0044 | 0.0064 | 0.002 |
| E369 | $M215^{5.54}$ | 0.0067 | 0.0036 | -0.0031 |
| Q370 | $M215^{5.54}$ | 0.0034 | 0.0038 | 0.0003 |
| S396 | $M215^{5.54}$ | 0.0017 | 0.0017 | 0 |
| S401 | $M215^{5.54}$ | 0.0076 | 0.0076 | 0 |
| E371 | $F217^{5.56}$ | 0.0039 | 0.0029 | -0.001 |
| E369 | $V218^{5.57}$ | 0.0002 | 0.0021 | 0.0019 |
| E371 | $V218^{5.57}$ | 0.0127 | 0.0079 | -0.0048 |
| E373 | $V218^{5.57}$ | 0.0061 | 0.0067 | 0.0006 |

| | | | | |
|---|---|---|---|---|
| N398 | V218$^{5.57}$ | 0.008 | 0.011 | 0.003 |
| S401 | Y219$^{5.58}$ | -0.013 | -0.0032 | 0.0098 |
| E369 | S220$^{5.59}$ | 0.0042 | 0.009 | 0.0048 |
| Q370 | S220$^{5.59}$ | 0.0046 | 0.0072 | 0.0026 |
| E371 | S220$^{5.59}$ | 0.0127 | 0.0136 | 0.0008 |
| T360 | M279$^{6.41}$ | 0.0081 | 0.0081 | 0 |
| S364 | M279$^{6.41}$ | 0.0087 | 0.0087 | 0 |
| S396 | M279$^{6.41}$ | 0.0015 | 0.0015 | 0 |
| S401 | M279$^{6.41}$ | 0.0011 | 0.0011 | 0 |
| S411 | M279$^{6.41}$ | 0.0011 | 0.0058 | 0.0046 |
| T360 | F282$^{6.44}$ | 0.008 | 0.008 | 0 |
| S364 | F282$^{6.44}$ | 0.0062 | 0.0062 | 0 |
| S396 | F282$^{6.44}$ | 0.0036 | 0.0036 | 0 |
| S411 | F282$^{6.44}$ | 0.0033 | 0.0033 | 0 |
| E369 | C285$^{6.47}$ | 0.0167 | 0.0093 | -0.0074 |
| E369 | E62 | 0.0081 | 0.0077 | -0.0004 |
| Q370 | R63 | 0.0141 | 0.0094 | -0.0046 |
| E371 | R63 | 0.0095 | 0.0079 | -0.0016 |
| E369 | V67$^{2.38}$ | 0.0251 | 0.0227 | -0.0024 |
| Q370 | T68$^{2.39}$ | 0.0175 | 0.0132 | -0.0043 |
| Q370 | R131$^{3.50}$ | 0.019 | 0.0094 | -0.0096 |
| E369 | Y132$^{3.51}$ | 0.0043 | 0.0042 | -0.0001 |
| Q370 | Y132$^{3.51}$ | 0.0041 | 0.003 | -0.0011 |
| E371 | Y132$^{3.51}$ | 0.0111 | 0.0103 | -0.0008 |
| R343 | P138 | -0.018 | -0.0053 | 0.0127 |
| Y354 | K147$^{4.39}$ | 0.0225 | 0.0225 | 0 |
| L342 | N148$^{4.40}$ | 0.0243 | 0.0132 | -0.011 |
| D410 | K149$^{4.41}$ | -0.0199 | -0.0098 | 0.0101 |
| N398 | A150$^{4.42}$ | 0.0065 | -0.0021 | -0.0086 |
| K348 | R151$^{4.43}$ | 0.0224 | 0.0145 | -0.0079 |
| Q370 | R151$^{4.43}$ | 0.0243 | 0.0125 | -0.0118 |
| N398 | R151$^{4.43}$ | 0.0067 | 0.0003 | -0.0065 |
| E369 | V152$^{4.44}$ | 0.0184 | 0.014 | -0.0043 |
| Q370 | V152$^{4.44}$ | 0.0184 | 0.0137 | -0.0046 |
| E369 | I153$^{4.45}$ | 0.0015 | 0.0003 | -0.0011 |
| S356 | I154$^{4.46}$ | 0.0217 | 0.0217 | 0 |
| Q363 | I154$^{4.46}$ | 0.0197 | 0.0197 | 0 |
| S364 | I154$^{4.46}$ | 0.0184 | 0.0184 | 0 |
| H390 | L230$^{5.69}$ | 0.0136 | 0.0136 | 0 |
| T408 | L230$^{5.69}$ | 0.0113 | 0.0075 | -0.0038 |
| S411 | E237 | -0.0163 | -0.0127 | 0.0037 |
| S346 | R239 | -0.0168 | -0.0106 | 0.0062 |
| G353 | R239 | 0.0017 | 0.0049 | 0.0032 |
| R344 | Q247 | -0.0171 | -0.0137 | 0.0035 |
| K348 | G252 | 0.0125 | 0.0047 | -0.0078 |
| S411 | H256 | -0.0184 | -0.0124 | 0.006 |

| | | | | |
|---|---|---|---|---|
| N409 | C265$^{6.27}$ | -0.0203 | -0.0085 | 0.0118 |
| E369 | K267$^{6.29}$ | 0.0166 | 0.0072 | -0.0094 |
| G392 | E268$^{6.30}$ | 0.0127 | 0.0127 | 0 |
| G383 | N322$^{7.49}$ | -0.0232 | -0.0082 | 0.015 |
| T384 | N322$^{7.49}$ | -0.0268 | -0.0077 | 0.0191 |
| P395 | N322$^{7.49}$ | -0.017 | -0.0017 | 0.0153 |
| S396 | N322$^{7.49}$ | -0.0149 | -0.0096 | 0.0053 |
| N352 | P323$^{7.50}$ | 0.0164 | 0.0131 | -0.0033 |
| T384 | P323$^{7.50}$ | -0.0164 | 0.0072 | 0.0236 |
| E385 | P323$^{7.50}$ | -0.022 | -0.0028 | 0.0192 |
| D386 | P323$^{7.50}$ | -0.0196 | 0.0025 | 0.0221 |
| N409 | P323$^{7.50}$ | -0.0294 | -0.0089 | 0.0205 |
| D410 | P323$^{7.50}$ | -0.0164 | 0.0002 | 0.0166 |
| D410 | L324$^{7.51}$ | -0.0198 | -0.0063 | 0.0134 |
| R344 | I325$^{7.52}$ | -0.0212 | -0.0127 | 0.0085 |
| D410 | I325$^{7.52}$ | -0.016 | -0.0147 | 0.0012 |
| L377 | Y326$^{7.53}$ | 0.0172 | 0.0172 | 0 |
| G353 | C327$^{7.54}$ | 0.014 | 0.0044 | -0.0096 |
| G353 | F332$^{8.50}$ | 0.0149 | 0.01 | -0.005 |
| G392 | F332$^{8.50}$ | 0.015 | 0.015 | 0 |
| A349 | R333$^{8.51}$ | 0.0142 | 0.0121 | -0.0021 |
| D386 | R333$^{8.51}$ | -0.0182 | -0.0042 | 0.014 |
| H390 | I334$^{8.52}$ | 0.017 | 0.017 | 0 |
| S346 | F336$^{8.54}$ | 0.0168 | 0.0091 | -0.0077 |
| G392 | Q337$^{8.55}$ | 0.016 | 0.016 | 0 |
| E373 | L339$^{8.57}$ | 0.0144 | 0.0144 | 0 |
| N322$^{7.49}$ | V48$^{1.47}$ | 0.0245 | 0.0087 | -0.0158 |
| C327$^{7.54}$ | D79$^{2.50}$ | -0.0225 | -0.0031 | 0.0194 |
| P323$^{7.50}$ | L80$^{2.51}$ | 0.0262 | 0.0055 | -0.0207 |
| L324$^{7.51}$ | L80$^{2.51}$ | 0.0228 | 0.0081 | -0.0147 |
| V152$^{4.44}$ | V81$^{2.52}$ | -0.02 | -0.0082 | 0.0119 |
| C327$^{7.54}$ | V81$^{2.52}$ | -0.0263 | -0.0051 | 0.0213 |
| F240 | T118$^{3.37}$ | 0.0237 | 0.0128 | -0.0109 |
| T66$^{2.37}$ | I121$^{3.40}$ | 0.0231 | 0.001 | -0.022 |
| N69$^{2.40}$ | L124$^{3.43}$ | 0.0262 | 0.0037 | -0.0225 |
| I154$^{4.46}$ | V157$^{4.49}$ | -0.0188 | -0.0132 | 0.0056 |
| V152$^{4.44}$ | V213$^{5.52}$ | -0.0235 | -0.0102 | 0.0134 |
| F336$^{8.54}$ | V213$^{5.52}$ | -0.0284 | -0.0206 | 0.0078 |
| N148$^{4.40}$ | I214$^{5.53}$ | 0.0137 | 0.0089 | -0.0048 |
| L230$^{5.69}$ | I214$^{5.53}$ | 0.0083 | 0.0093 | 0.0011 |
| K235$^{5.74}$ | I214$^{5.53}$ | 0.0108 | 0.0125 | 0.0018 |
| F264$^{6.26}$ | I214$^{5.53}$ | -0.0126 | -0.0029 | 0.0097 |
| C265$^{6.27}$ | I214$^{5.53}$ | 0.0135 | 0.0072 | -0.0062 |
| R131$^{3.50}$ | M215$^{5.54}$ | 0.0102 | 0.0044 | -0.0057 |
| R228$^{5.67}$ | M215$^{5.54}$ | 0.0083 | 0.0009 | -0.0075 |
| K232$^{5.71}$ | M215$^{5.54}$ | 0.0118 | 0.0105 | -0.0013 |

| | | | | |
|---|---|---|---|---|
| $D234^{5.73}$ | $M215^{5.54}$ | 0.0131 | 0.014 | 0.001 |
| $K235^{5.74}$ | $M215^{5.54}$ | 0.0081 | 0.0094 | 0.0013 |
| $S262^{6.24}$ | $M215^{5.54}$ | 0.0133 | 0.0101 | -0.0032 |
| $K263^{6.25}$ | $M215^{5.54}$ | 0.0115 | 0.0102 | -0.0013 |
| $C265^{6.27}$ | $M215^{5.54}$ | 0.0149 | 0.0107 | -0.0042 |
| $K232^{5.71}$ | $V216^{5.55}$ | 0.0104 | 0.0094 | -0.001 |
| $I233^{5.72}$ | $V216^{5.55}$ | 0.0163 | 0.0138 | -0.0025 |
| $I135^{3.54}$ | $F217^{5.56}$ | -0.0198 | -0.0189 | 0.0009 |
| $V152^{4.44}$ | $F217^{5.56}$ | -0.0266 | -0.0189 | 0.0077 |
| $E268^{6.30}$ | $F217^{5.56}$ | -0.0142 | -0.0094 | 0.0048 |
| $R228^{5.67}$ | $V218^{5.57}$ | 0.0108 | 0.0089 | -0.0019 |
| $K232^{5.71}$ | $V218^{5.57}$ | 0.0111 | 0.0086 | -0.0025 |
| F240 | $V218^{5.57}$ | 0.0244 | 0.0092 | -0.0153 |
| $C265^{6.27}$ | $V218^{5.57}$ | 0.0116 | 0.0072 | -0.0044 |
| $R131^{3.50}$ | $Y219^{5.58}$ | -0.0214 | -0.0206 | 0.0008 |
| L145 | $Y219^{5.58}$ | -0.0237 | -0.0078 | 0.0158 |
| $R151^{4.43}$ | $Y219^{5.58}$ | -0.0187 | -0.0129 | 0.0058 |
| $V152^{4.44}$ | $Y219^{5.58}$ | -0.0206 | -0.0148 | 0.0058 |
| $I154^{4.46}$ | $Y219^{5.58}$ | -0.0184 | -0.0111 | 0.0073 |
| $K235^{5.74}$ | $Y219^{5.58}$ | 0.0151 | 0.0114 | -0.0038 |
| E62 | $Y219^{5.58}$ | -0.0249 | -0.0185 | 0.0064 |
| L64 | $Y219^{5.58}$ | -0.0227 | -0.0152 | 0.0076 |
| $V67^{2.38}$ | $Y219^{5.58}$ | -0.0174 | -0.0105 | 0.0069 |
| $K232^{5.71}$ | $M279^{6.41}$ | 0.0123 | 0.0097 | -0.0026 |
| $C265^{6.27}$ | $M279^{6.41}$ | 0.0138 | 0.0064 | -0.0074 |
| $K232^{5.71}$ | $G280^{6.42}$ | 0.0091 | 0.0098 | 0.0007 |
| $I233^{5.72}$ | $G280^{6.42}$ | 0.0126 | 0.0165 | 0.0039 |
| $D234^{5.73}$ | $G280^{6.42}$ | 0.0126 | 0.0067 | -0.0059 |
| $K263^{6.25}$ | $G280^{6.42}$ | 0.0083 | 0.0103 | 0.002 |
| $F264^{6.26}$ | $T281^{6.43}$ | -0.012 | -0.0018 | 0.0102 |
| $K267^{6.29}$ | $T281^{6.43}$ | 0.0107 | 0.0028 | -0.008 |
| $R228^{5.67}$ | $F282^{6.44}$ | 0.0108 | 0.004 | -0.0068 |
| $Q231^{5.70}$ | $F282^{6.44}$ | 0.0088 | 0.0078 | -0.001 |
| $K232^{5.71}$ | $F282^{6.44}$ | 0.0122 | 0.0074 | -0.0048 |
| $I233^{5.72}$ | $F282^{6.44}$ | 0.0134 | 0.0074 | -0.006 |
| $D234^{5.73}$ | $F282^{6.44}$ | 0.0186 | 0.0102 | -0.0084 |
| $K235^{5.74}$ | $F282^{6.44}$ | 0.0099 | 0.0081 | -0.0018 |
| $C265^{6.27}$ | $F282^{6.44}$ | 0.011 | 0.0017 | -0.0093 |
| $R228^{5.67}$ | $T283^{6.45}$ | 0.0086 | 0.0046 | -0.004 |
| $L230^{5.69}$ | $T283^{6.45}$ | -0.0107 | -0.0002 | 0.0105 |
| $Q231^{5.70}$ | $T283^{6.45}$ | 0.0082 | 0.0071 | -0.0011 |
| $K235^{5.74}$ | $T283^{6.45}$ | 0.0114 | 0.004 | -0.0074 |
| $C265^{6.27}$ | $T283^{6.45}$ | 0.0084 | 0.0063 | -0.0022 |
| $L266^{6.28}$ | $T283^{6.45}$ | 0.0121 | 0.0049 | -0.0072 |
| $R151^{4.43}$ | $L284^{6.46}$ | -0.0177 | -0.0131 | 0.0045 |
| $E268^{6.30}$ | $L284^{6.46}$ | -0.0109 | -0.0031 | 0.0078 |

| | | | | |
|---|---|---|---|---|
| H269$^{6.31}$ | L284$^{6.46}$ | -0.0119 | -0.0131 | -0.0012 |
| R151$^{4.43}$ | S319$^{7.46}$ | -0.0221 | -0.006 | 0.0161 |
| V152$^{4.44}$ | S319$^{7.46}$ | -0.0228 | -0.0124 | 0.0104 |
| I154$^{4.46}$ | S319$^{7.46}$ | -0.0226 | -0.0084 | 0.0142 |
| L64 | S319$^{7.46}$ | -0.0208 | -0.0086 | 0.0122 |
